# Unique epigenomic signatures identify biologically significant subtypes of MDS and predict response to azacitidine

**DOI:** 10.1101/2025.04.23.650336

**Authors:** Qin Yang, Miguel Torres-Martin, Masataka Taguchi, Alice Brogi, Irene Casalin, Eleonora Ceneri, Stephanie Halene, Amy E. DeZern, Elizabeth A. Griffiths, Matilde Y. Follo, Carlo Finelli, Jerald P. Radich, Michael J. Rauh, Rafael Bejar, Mikkael A. Sekeres, Valeria Santini, Maria E. Figueroa

**Affiliations:** Dept. of Biochemistry & Molecular Biology, University of Miami Miller School of Medicine, Miami, FL, USA; Sylvester Comprehensive Cancer Center, University of Miami Miller School of Medicine, Miami, FL, USA; MDS Unit, Hematology, DMSC University of Florence, AOU Careggi, Florence, Italy; Dept. of Biomedical and Neuromotor Sciences, University of Bologna, Bologna, Italy; Dept. of Internal Medicine, Yale University School of Medicine, New Haven, CT, USA; Sidney Kimmel Comprehensive Cancer Center, Johns Hopkins Medicine, Baltimore, MD, United States; Roswell Park Comprehensive Cancer Center, Buffalo, NY, USA; Dept. of Experimental, Diagnostic and Specialty Medicine, Institute of Hematology L. and A. Seràgnoli, University of Bologna, S. Orsola-Malpighi Hospital, Bologna, Italy; Clinical Research Division, Fred Hutchinson Cancer Center, Seattle, WA, USA; Department of Pathology and Molecular Medicine, Queen’s University, Kingston, Canada; Division of Hematology and Oncology, University of California San Diego Moores Cancer Center, La Jolla, CA, USA; Div. of Hematology, University of Miami Miller School of Medicine, Miami, FL, USA; Dept. of Human Genetics, University of Miami Miller School of Medicine, Miami, FL, USA

**Keywords:** Myelodysplastic syndromes, DNA methylation, epigenetics, response classifiers, hypomethylating agents, azacitidine

## Abstract

Myelodysplastic syndromes (MDS) are characterized by aberrant DNA methylation, and mutations in epigenetic modifiers are frequently found in these patients. Although DNA methyltransferase inhibitors (DNMTi) are used to treat MDS, response variability remains a challenge in the clinic, with limited predictive markers. Through comprehensive genomic, epigenomic, and transcriptomic analyses, we have gained valuable insights into the intricate interplay between genetic and epigenetic alterations in MDS. We describe aberrantly hyper and hypomethylated regions in MDS, extending beyond promoter regions and affecting long-distance regulatory elements. Using these aberrant DNA methylation patterns, we classified MDS patients into epigenetic subtypes correlated with known molecular drivers. This epigenetic classification includes a novel group of patients characterized only by their shared DNA methylation profile and lacking any genetic drivers. Furthermore, we identified a robust DNA methylation signature capable of distinguishing DNMTi responders from non-responders prior to receiving treatment. Leveraging these DMRs, we developed robust classifiers capable of predictive response to DNMTi by integrating DNA methylation, gene expression, mutations, and laboratory parameters. Our findings highlight the potential of epigenetic-based classifiers for personalized treatment approaches for MDS patients.

**Key Points:** DNA methylation patterns define biologically meaningful MDS subtypes and uncover a new group lacking known mutations.

A methylation-based signature at diagnosis predicts azacitidine response, supporting its use in guiding personalized MDS therapy.

## INTRODUCTION

Myelodysplastic syndromes (MDS) encompass a heterogeneous group of disorders characterized by ineffective hematopoiesis, dysplastic changes in the bone marrow (BM), and the potential to progress to acute myeloid leukemia (AML)^2–4^. Aberrant DNA methylation (DNAme) is a hallmark of MDS; however, its role remains incompletely understood. Early reports focused on aberrant hypermethylation of tumor suppressor gene promoters, e.g., p15, E-cadherin^5–7^, while the advent of DNAme microarrays revealed widespread aberrant hypermethylation at diagnosis^8^, which directly correlated with disease aggressiveness and progression^9^. Later molecular studies reported the existence of recurrent mutations in genes involved in epigenetic regulation, such as *TET2, ASXL1, DNMT3A*, and *EZH2* in MDS^10–12^. However, these groundbreaking early studies used smaller cohorts from single centers, no CD34+ selection, and analysis restricted to promoter methylation and gene expression microarrays.

Unlike de novo AML, MDS is more likely to be resistant to conventional chemotherapy^13,14^. For higher-risk MDS patients, while hematopoietic cell transplantation (HCT) offers the only curative option, it is usually available to only a small fraction of this older patient population. The remaining higher-risk patients rely on treatment with DNMTi – azacitidine (AZA) or decitabine (DAC)– the only available treatment option that has demonstrated a survival advantage compared to conventional care^15–19^. However, the overall response rate to these agents is between 30-40%, responses may take longer than six months^18^, and the duration lasts a median of 17 months. Consequently, for the majority of patients in real-world settings, these agents are ineffective, expose them to side effects, and result in significant time and economic burden^15^. Once DNMTi have failed, survival in the absence of an allogeneic HCT is generally less than six months^20^. It is thus critical to identify MDS patients who are unlikely to benefit from DNMTi to accelerate access to HCT, offer up-front investigational studies, and, for frailer patients, offer the option of supportive care alone.

Despite some studies suggesting associations between the methylation reversal at select loci and clinical response to DNMTis^21–24^, overall, prior epigenetic studies have failed to identify robust associations between specific baseline DNAme profiles and DNMTi response^21,23,25,26^. The absence of response classifiers impedes the customization of treatment plans for individual MDS patients. To address this critical gap, we conducted a multicenter study of a cohort of AZA-treated MDS patients. Through genomic, epigenomic, and transcriptomic analyses on CD34^+^ BM cells, we identified widespread aberrant DNAme, targeting distal regulatory elements at key biological pathways. These aberrant epigenomic patterns can be used to classify MDS into distinct epigenetic subtypes linked to specific genetic, molecular, and clinical characteristics. Furthermore, we harnessed these epigenetic profiles to develop robust, epigenetic-based classifiers that correlate with accurate pre-treatment prediction of therapeutic response to AZA. These classifiers offer the potential to improve personalized treatment strategies and enhance outcomes for patients with MDS.

## METHODS

### MDS samples

Clinical BM specimens were collected from 98 de-identified MDS specimens classified as Intermediate or higher-risk MDS by the International Prognostic Scoring System (IPSS)^27^. Mononuclear cells (MNC) were isolated through Ficoll density centrifugation at diagnosis and frozen for later use. Institutional Review Board (IRB) approval was obtained at the University of Miami Miller School of Medicine, Yale University School of Medicine, Roswell Park Cancer Institute, University of Florence, University of Bologna, and the Southwestern Oncology Group (SWOG) central IRB and local IRBs of participating institutions. Written informed consent was obtained from all patients at the time of collection in accordance with the Declaration of Helsinki.

CD34⁺ cells were isolated using MACS magnetic beads. DNA and RNA were extracted with the QIAGEN AllPrep kit. Further details are provided in the Supplementary Methods.

### Statistical analysis

All statistical analyses were performed using R statistical software^28^ (version 4.1, R-Core Team, http://www.R-project.org/). For sequencing-based assays, significance was defined as q ≤ 0.05; for clinical comparisons, p ≤ 0.05 was used. Significance levels are denoted in figures as follows: *p ≤ 0.05, **p ≤ 0.01, ***p ≤ 0.001. Additional statistical details are provided in the Supplementary Methods.

### Software availability

Code used for the analyses in this study is available on Github: https://github.com/santur90/MDS_AZA

## RESULTS

### MDS displays extensive aberrant DNA hypomethylation affecting distal regulatory elements

To assess epigenetic deregulation in MDS, we analyzed pre-treatment CD34+ BM specimens from a multicenter cohort of 98 IPSS intermediate- or higher-risk MDS patients.^27^ The cohort consisted of cases from the North American Intergroup MDS trial S1117 (n=37 cases), augmented with cases from US (n=24) and Italian (n=37) academic centers. All patients received AZA-based regimens and had documented responses. Patients were studied with targeted gene mutation profiling of 112 genes frequently mutated in MDS (n=92 patients), DNAme by enhanced reduced representation bisulfite sequencing (ERRBS; n=90 cases), and gene expression by RNA-seq (n=77 cases) (**Table 1, Supplemental Table 1)**. We first compared gene expression profiles between CD34+ hematopoietic stem and progenitor cells (HSPC) from MDS patients and age-matched healthy individuals (n=14). Unsupervised analysis using principal components (PC) revealed modest discrimination of the two groups **(Figure 1A, left**). This was confirmed by orthogonal approaches using hierarchical clustering and correspondence analysis **(Supplemental Figure 1A)**. Furthermore, a direct comparison identified only 53 differentially expressed genes (DEGs; absolute fold change (FC) ≥ 1.5 and a false discovery rate (FDR) ≤ 0.05; **Supplemental Table 2**) between MDS and controls **(Supplemental Figure 1B)**. Gene set enrichment analysis (GSEA) identified upregulation of pathways related to interferon responses, IL6 signaling, oxidative phosphorylation, reactive oxygen species (ROS), cell cycle, and mitophagy in MDS **(Supplemental Figure 1C, Supplemental Table 3)**.

**Figure 1.**
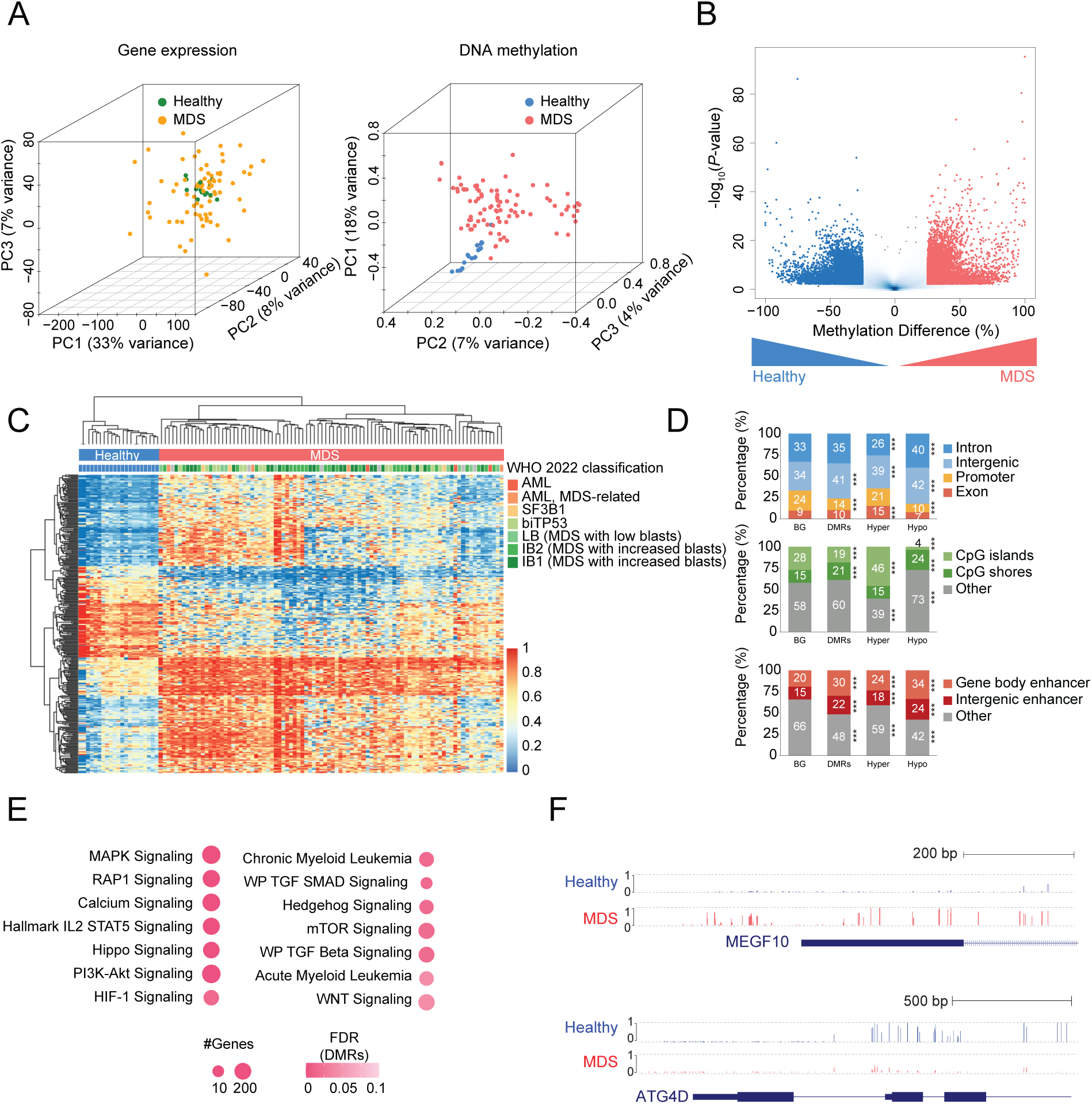
Comprehensive analysis of DNA methylation alterations in MDS. **A:** Principal Component Analysis of RNA-seq data (*left)* from healthy controls (green) and MDS patients (yellow), along with Enhanced Reduced Representation Bisulfite Sequencing *(right)* data from healthy controls (blue) and MDS patients (red). **B:** Volcano plot depicting methylation differences versus the -log_10_(*P*-value) between healthy controls and MDS patients. Hypomethylated and hypermethylated regions in MDS are shown in blue and red, respectively (absolute methylation difference ≥ 25%, FDR ≤ 0.05). **C:** Heatmap with hierarchical clustering illustrating methylation levels of differentially methylated regions (without missing values) in healthy controls (blue bar) and MDS patients (red bar), annotated with WHO 2022 subtypes. **D:** Stacked bar plot showing the distribution of differentially methylated regions across genomic regions, CpG islands, and enhancers. *P* values were calculated using a binomial test. ***, *P* ≤ 0.001. **E:** Bubble plot depicting enriched pathways (FDR ≤ 0.1) for MDS-associated differentially methylated regions, using all CpG tiles as the background. Bubble size corresponds to the number of genes in each pathway, color indicates statistical significance. **F:** UCSC Genome Browser tracks of two representative loci illustrating hypermethylated (*top*) and hypomethylated (*bottom*) regions in MDS. Bar height represents the methylation level of each region.

**Table 1:**
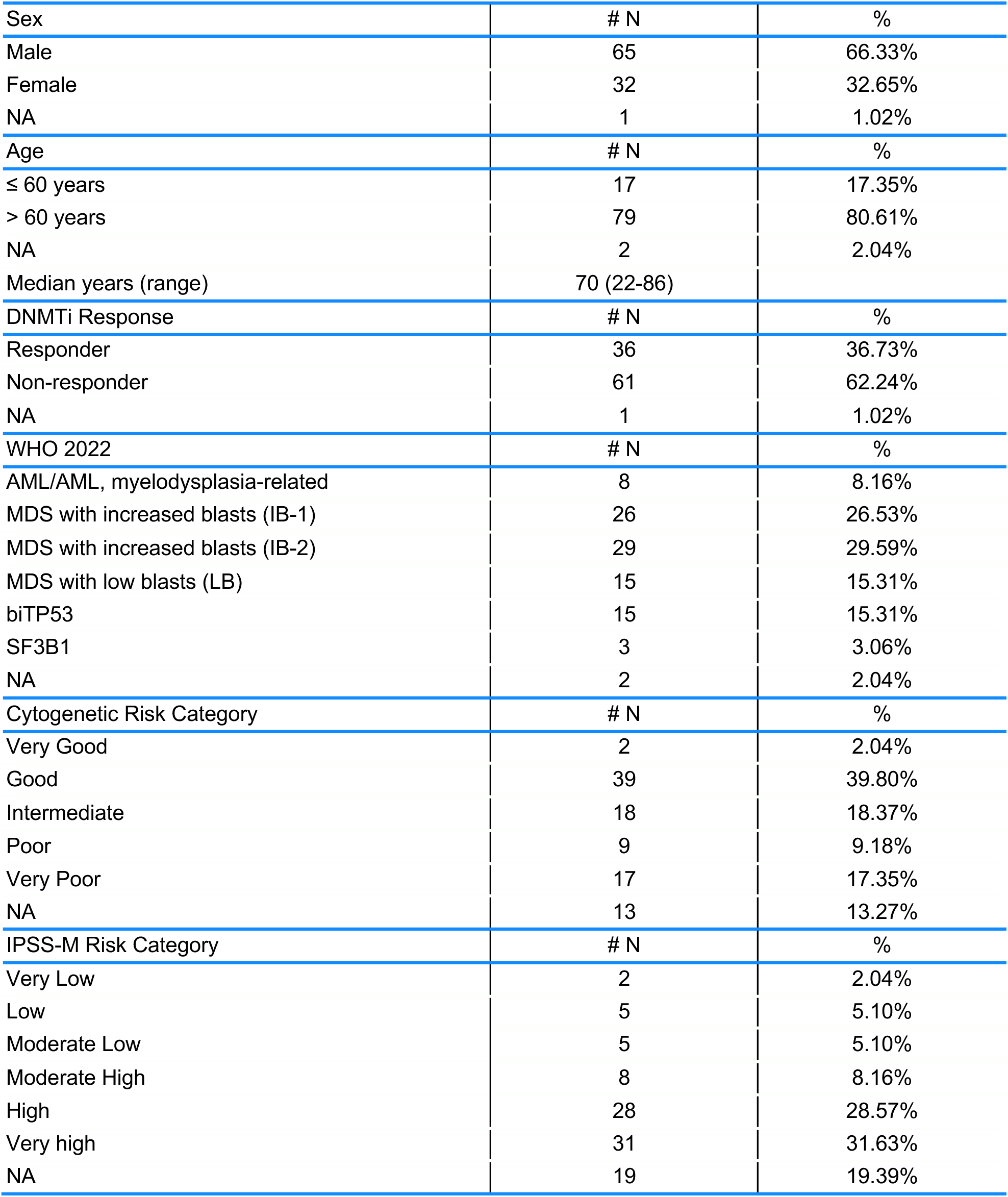
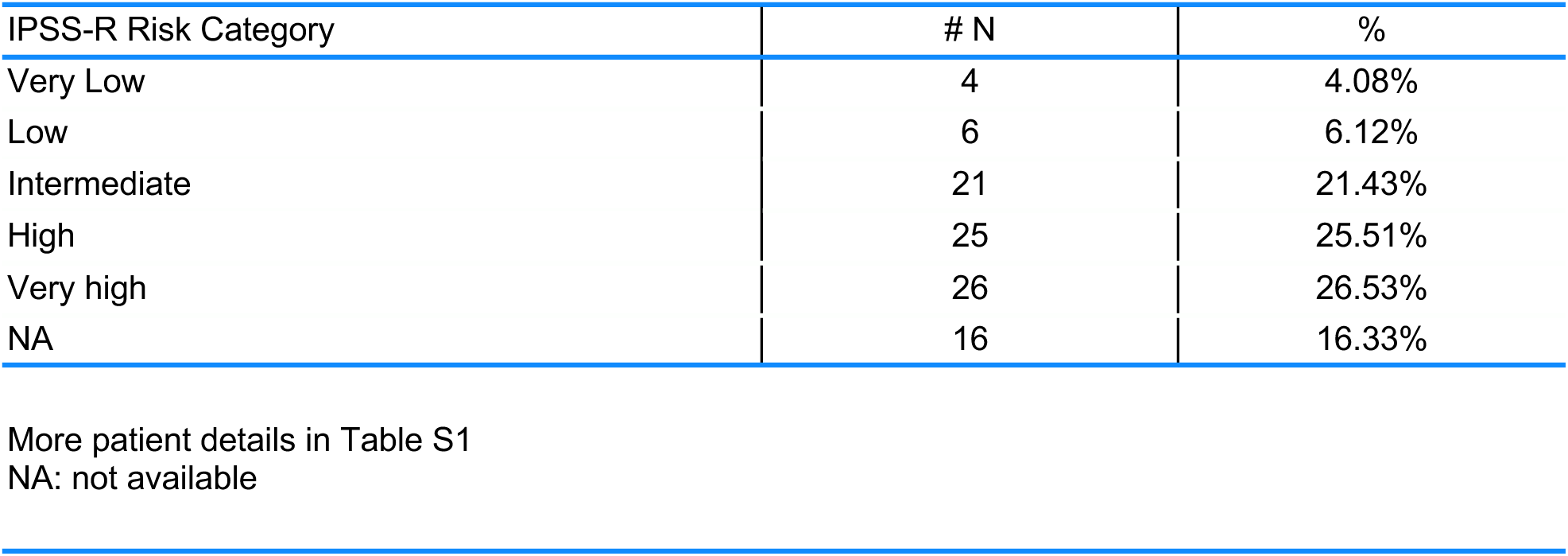
Patient Characteristics.

Next, we investigated DNAme patterns in the same cohort of MDS patients and samples from 21 healthy individuals^29^ using ERRBS. PC analysis based on DNAme profiles demonstrated a clear separation between healthy individuals and MDS patients **(Figure 1A, right)**. Notably, unlike prior studies focusing solely on promoter regions that captured only aberrant DNA hypermethylation in MDS^8,9^, supervised analysis comparing the two groups identified almost twice as many hypomethylated regions (n=15,398 differentially methylated regions (DMRs); absolute methylation difference ≤ 25% and FDR ≤ 0.05) in MDS than hypermethylated ones (n= 8,805) (**Figure 1B-C; Supplemental Table 4)**.

Since ERRBS targets not only promoters but also intronic, exonic, and distal intergenic regions, we were able to explore the impact of aberrant DNAme within specific genomic features. Overall, DMRs were enriched at intergenic regions (DMRs 41% vs. background 34%, binomial test p-value ≤0.001) while being less likely to target promoter regions (DMRs 14% vs. 24%, p-value ≤0.001). Hypermethylated DMRs were depleted from intronic regions (hyper-DMRs 26% vs. 33%, p-value ≤0.001), whereas hypomethylated DMRs displayed the opposite trend (hypo-DMRs 40% vs. 33%, p-value ≤0.001; **Figure 1D, *top***). Furthermore, we observed a significant enrichment of hypermethylated DMRs at CpG islands (hyper-DMRs 46% vs. 28%, p-value ≤0.001) while hypomethylated DMRs were enriched at CpG island shores (hypo-DMRs 24% vs. 15%, p-value ≤0.001; **Figure 1D, *middle***). Notably, both hypermethylated and hypomethylated DMRs were significantly enrichment at both gene body enhancers (DMRs 30% vs. 20%, p-value ≤ 0.001) and intergenic enhancers (DMRs 22% vs. 15%, p-value ≤0.001; **Figure 1D, *bottom***). Pathway enrichment analysis annotating these DMRs to neighboring transcripts showed enrichment in various leukemia pathways and signaling pathways known to play a role in hematopoiesis and malignant transformation, such as the Hippo^28,30^, WNT^31^, RAP1^32,33^, mTOR^34,35^, and MAPK^36,37^ signaling pathways. **(Figure 1E-F; Supplemental Table 3)**.

Taken together, these findings underscore the magnitude of aberrant gains and losses in DNAme in MDS beyond promoter regions, identifying crucial DNAme changes targeting distal regulatory elements. Furthermore, they provide insight into the potential functional implications of DNAme alterations in MDS, suggesting their involvement in critical signaling cascades and molecular processes known to regulate HSPC biology.

### DNAme profiles capture the underlying molecular and clinical heterogeneity of the disease

We hypothesized that DNAme profiles could serve to stratify MDS patients into distinct epigenetic subtypes reflective of the disease’s molecular and biological heterogeneity. Using hierarchical clustering, we classified MDS patients in our cohort into seven clusters and analyzed each cluster with respect to associated clinical features (age, sex), laboratory parameters (complete blood counts, blast percentages), molecular and cytogenetic information, WHO 2022 classification and AZA response. Specific molecular or cytogenetic features were considered cluster-defining when they showed both statistically significant enrichment and were associated with more than 40% of patients in a given cluster. **Table 2** describes the clinical, cytogenetic, and classification details for each of the 7 clusters.

**Table 2:** Summary of Patient Characteristics within each of the 7 DNA methylation Clusters.

| Cluster | I | II | III | IV | V | VI | VII | Others | # N |
| --- | --- | --- | --- | --- | --- | --- | --- | --- | --- |
| # Patient number | 13 | 13 | 6 | 6 | 19 | 14 | 14 | 5 | 90 |
| Sex |  |  |  |  |  |  |  |  |  |
| Male | 7 | 9 | 4 | 1 | **16\*** | 12 | 9 | 2 | 60 |
| Female | 6 | 4 | 2 | **5\*** | 3 | 2 | 5 | 3 | 30 |
| Age |  |  |  |  |  |  |  |  |  |
| ≤ 60 years | 0 | 4 | 1 | 1 | 5 | 2 | 3 | 0 | 16 |
| > 60 years | **13\*** | 9 | 5 | 5 | 14 | 12 | 10 | 5 | 73 |
| Median years (range) | 78<br>(69-85) | 67<br>(56-82) | 67<br>(54-86) | 76<br>(48-85) | 67<br>(47-84) | 70<br>(48-81) | 72<br>(37-84) | 76<br>(62-82) |  |
| NA | 0 | 0 | 0 | 0 | 0 | 0 | 1 | 0 | 1 |
| DNMTi Response |  |  |  |  |  |  |  |  |  |
| Responder | 3 | 4 | 1 | 2 | 8 | **10\*\*** | 5 | 0 | 33 |
| Non-responder | 10 | 9 | 5 | 4 | 11 | 4 | 9 | 5 | 57 |
| WHO 2022 |  |  |  |  |  |  |  |  |  |
| AML/AML,<br>myelodysplasia-related | 2 | 1 | 0 | 0 | 0 | 2 | 2 | 0 | 7 |
| MDS with increased<br>blasts (IB-1) | 2 | 6 | 4 | 2 | 6 | 2 | 2 | 1 | 25 |
| MDS with increased<br>blasts (IB-2) | 7 | 5 | 2 | 1 | 3 | 5 | 3 | 1 | 27 |
| MDS with low blasts (LB) | 1 | 1 | 0 | 0 | 1 | 4 | 3 | 3 | 13 |
| biTP53 | 1 | 0 | 0 | 1 | **9\*\*** | 1 | 2 | 0 | 14 |
| SF3B1 | 0 | 0 | 0 | **2\*** | 0 | 0 | 1 | 0 | 3 |
| NA | 0 | 0 | 0 | 0 | 0 | 0 | 1 | 0 | 1 |
| Cytogenetic Risk<br>Category |  |  |  |  |  |  |  |  |  |
| Very Good | 0 | 0 | 0 | 1 | 0 | 1 | 0 | 0 | 2 |
| Good | 5 | 9 | 3 | 1 | 4 | 7 | 6 | 3 | 38 |
| Intermediate | **6\*** | 2 | 1 | 1 | 2 | 1 | 4 | 0 | 17 |
| Poor | 0 | 1 | 0 | 1 | 3 | 1 | 0 | 1 | 7 |
| Very Poor | 2 | 0 | 0 | 1 | 7 | 1 | 4 | 1 | 16 |
| NA | 0 | 1 | 2 | 1 | 3 | 3 | 0 | 0 | 10 |
| IPSS-M Risk Category |  |  |  |  |  |  |  |  |  |
| Very Low | 0 | 0 | 0 | 0 | 0 | **2\*** | 0 | 0 | 2 |
| Low | 0 | 0 | 0 | 1 | 0 | 2 | 1 | 1 | 5 |
| Moderate Low | 0 | 0 | 0 | 0 | 0 | 3 | 2 | 0 | 5 |
| Moderate High | 1 | 3 | 0 | 0 | 2 | 0 | 2 | 0 | 8 |
| High | 7 | 3 | 3 | 1 | 5 | 3 | 3 | 2 | 27 |
| Very high | 5 | 5 | 1 | 3 | 9 | 1 | 4 | 2 | 30 |
| NA | 0 | 2 | 2 | 1 | 3 | 3 | 2 | 0 | 13 |

| IPSS-R Risk Category |  |  |  |  |  |  |  |  |  |
| --- | --- | --- | --- | --- | --- | --- | --- | --- | --- |
| Very Low | 0 | 0 | 0 | 1 | 0 | 2 | 0 | 1 | 4 |
| Low | 0 | 2 | 1 | 0 | 2 | 0 | 1 | 0 | 6 |
| Intermediate | 2 | 5 | 0 | 0 | 3 | 4 | 5 | 1 | 20 |
| High | 6 | 2 | 3 | 3 | 4 | 3 | 1 | 2 | 24 |
| Very high | 5 | 2 | 0 | 1 | 7 | 2 | 6 | 1 | 24 |
| NA | 0 | 2 | 2 | 1 | 3 | 3 | 1 | 0 | 12 |

Fisher's exact test: p-value≤0.05 (\*); p-value≤0.01 (\*\*)

As expected, mutations in splicing factor proteins and epigenetic modifiers were very frequently seen in the cohort, and most clusters included patients carrying some of these mutations. However, similar to what we had previously observed in the context of AML^38^, the DNAme cluster identity correlated strongly with specific clinical and molecular features **(Figure 2A-B)**. Cluster II (n=13) was enriched for patients carrying ASXL1 (Fisher’s test p-value≤0.01), STAG2 (p-value≤0.05), and RUNX1 (p-value≤0.05) mutations. Cluster III (n=6) was defined by the presence of U2AF1 (p-value≤0.01) and BCOR (p-value≤0.01) mutations. Cluster IV (n=6) was enriched for the presence of SF3B1 mutations (p-value≤0.001) co-occurring with RUNX1 mutations in female patients (p-value≤0.05). In contrast, 5/6 patients with SF3B1 outside of Cluster IV were male and failed to cluster together. Notably, 10/19 patients in Cluster V carried mutations in TP53 (p-value≤0.001). In addition, patients in this cluster were almost devoid of splicing factor mutations, with the few positive cases carrying almost exclusively U2AF1 mutations. Moreover, Cluster V was strongly enriched in male patients, with 16/19 male cases (p-value≤0.05). Notably, the remaining 8 cases with TP53 mutations were scattered among the other clusters and were mainly female (5/8). Cluster VII was enriched in mutations in the DNAme machinery (8/14), most of them (6/8) affecting DNMT3A (p-value≤0.05) (**Supplemental Table 5**).

**Figure 2.**
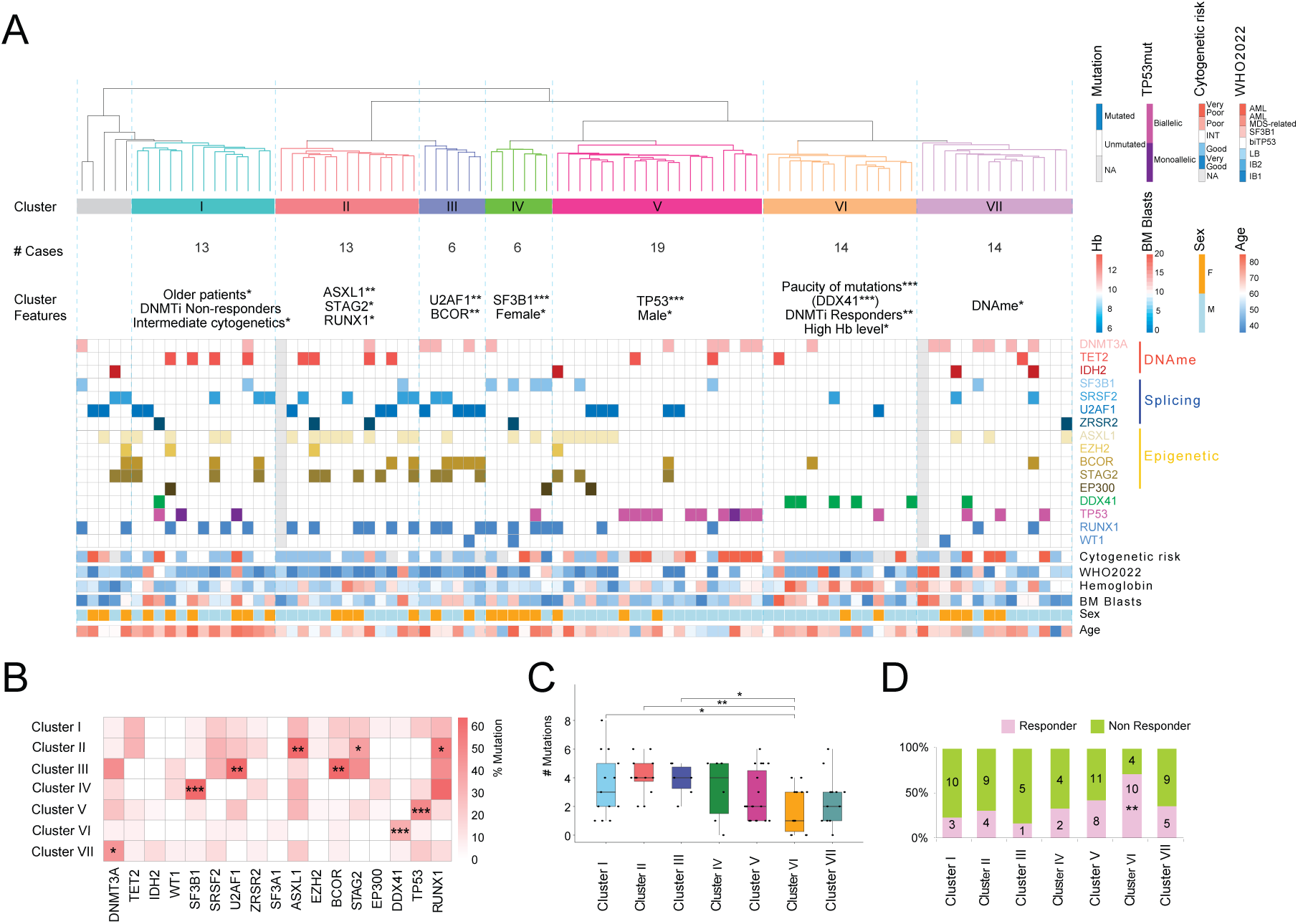
DNA methylation captures molecular heterogeneity in MDS. **A:** Heatmap with hierarchical clustering illustrating the seven MDS patient clusters identified based on DNA methylation data, with corresponding mutation and clinical annotations displayed below. Patients harboring gene mutations are color-coded. **B:** Heatmap depicting the mutation profiles of each identified cluster. *P* values were calculated using Fisher’s exact test. *, *P* ≤ 0.05; **, *P* ≤ 0.01; ***, *P* ≤ 0.001. **C:** Box plot showing the number of mutations per patient across the identified clusters. *P* values were calculated using the Wilcoxon rank sum test. *, *P* ≤ 0.05; **, *P* ≤ 0.01. **D:** Bar plot displaying the number of clinical responders and non-responders within each cluster. *P* values were calculated using Fisher’s exact test. **, *P* ≤ 0.01.

Two of the clusters identified could not be classified based on any specific mutation, but instead, patients in those clusters shared robust epigenetic profiles and clinical features. Patients in Cluster I (n=13) were characterized by older age (median 78 years old, range=69-85 vs. median 68, range = 37-86; p-value≤0.05) and intermediate-risk cytogenetics (p-value≤0.05), the majority of whom (n=10) did not respond to DNMTi treatment, though this did not achieve statistical significance.

Finally, unlike patients in Cluster II, all of whom carried at least one mutation (with most patients carrying three or more), Cluster VI was striking because of its much lower mutation frequency than other clusters **(Figure 2C)**. However, despite this overall paucity of mutations, it was noteworthy that 5 out of 7 patients harboring DDX41 mutations (4 germline and 1 somatic) in our cohort were located in this cluster p-value≤0.001). Despite this, DDX41 mutations were not considered cluster-defining as they affected only 5/14 patients (36%) in the cluster. In addition to a low frequency of mutations, patients in cluster VI showed a higher response rate to AZA than patients in other clusters (defined as a response of hematologic improvement, partial or complete response within 6 months of beginning treatment), with 71% (10 out of 14) responding to the drug (p-value≤0.01; **Figure 2D)**. In line with this paucity of mutations, 7 out of 12 patients from our MDS cohort that fell under the "Very Low," "Low," or "Moderate Low" 2022 IPSS-M risk categories were classified together in this cluster (p-value≤0.01) **(Supplemental Table 5)**.

Notably, and in line with our PC analysis findings, a comparable hierarchical clustering analysis based on gene expression data failed to classify patients into robust clusters correlated with molecular and clinical features (**Supplemental Figure 2**). Taken together, these findings highlight the robustness of DNAme in capturing the molecular and clinical heterogeneity of MDS.

### Cluster VI patients have mild epigenetic deregulation and shared clinical and laboratory features

To further characterize the epigenetic deregulation of the epigenetic clusters, we compared them to healthy controls. We first conducted a PC analysis to visualize the epigenetic distances between controls and MDS patient clusters **(Figure 3A).** Notably, cluster VI, which carried very few mutations and responded well to AZA treatment, was epigenetically closer to the healthy controls than any of the other clusters, indicating the DNAme profiles of these patients are less perturbed. Supervised analysis comparing each cluster to the controls revealed that cluster VI displayed a lower number of DMRs than the other clusters. While the other six clusters all displayed widespread epigenetic abnormalities that clearly distinguished them from controls, with total DMRs ranging from 22,000 to 71,000, Cluster VI had only 5,855 hypomethylated DMRs and 1,654 hypermethylated DMRs (absolute methylation difference ≥ 25% and FDR ≤ 0.05; **Figure 3B, Supplemental Figure 3 & 4 and Supplemental Table 6**). Furthermore, Cluster VI exhibited significantly higher levels of hemoglobin (Hb) compared to the other clusters (Mean: 10.9 g/dL; range: 7.7-13.7 g/dL; ANOVA p-value≤0.05), once again indicating that patients in this cluster display more benign features than the rest of the cohort. Notably, there were no significant differences in BM blast percentage, red blood cells, platelets, white blood cells, or absolute neutrophil counts between Cluster VI and the other clusters (**Figure 3C).**

**Figure 3.**
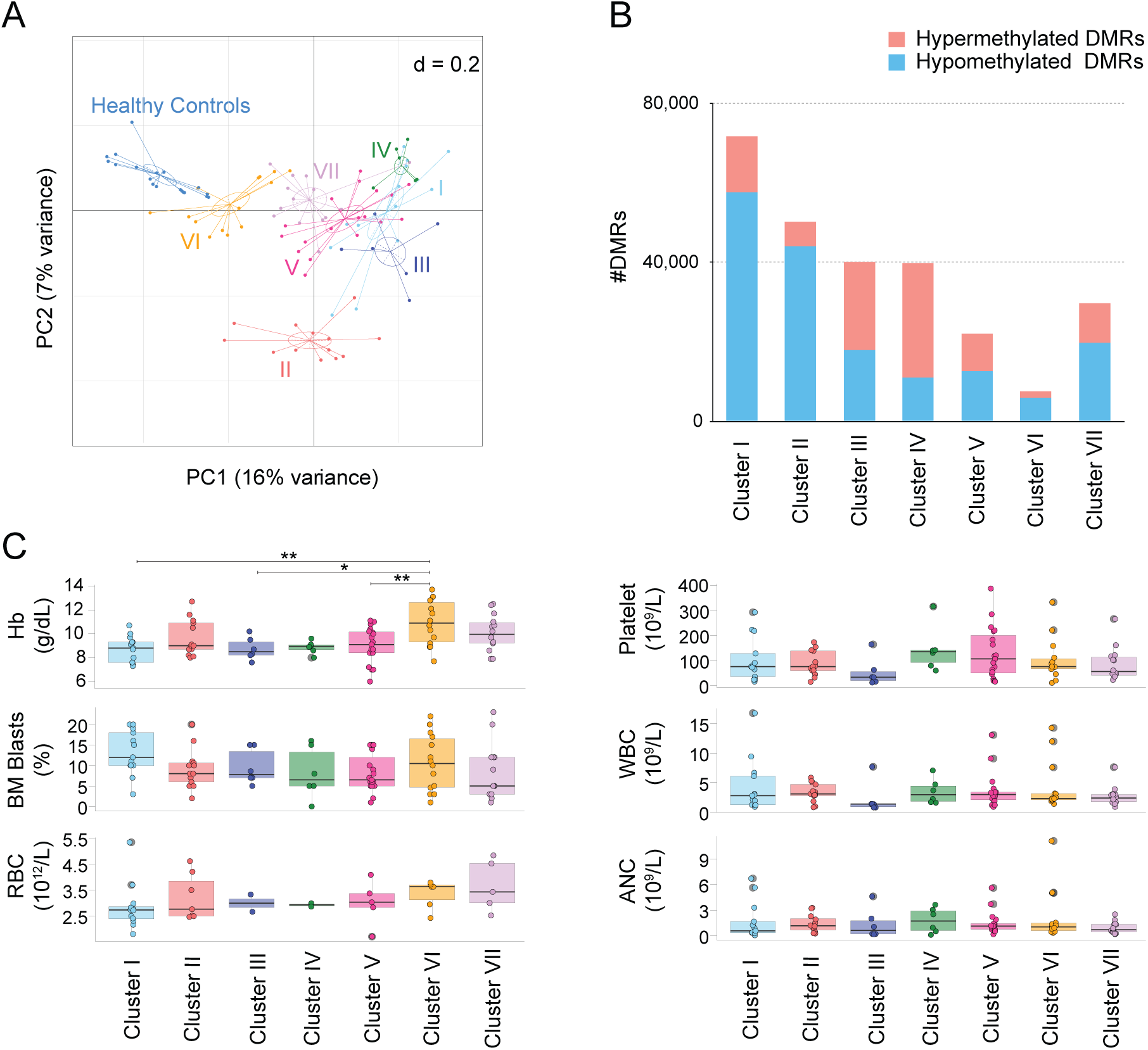
Epigenetic and clinical characteristics associated with each DNA methylation cluster. **A:** Principal Components Analysis of healthy controls (blue) and MDS patient clusters. **B:** Bar plot showing number of hypermethylated (red) and hypomethylated (blue) regions in each MDS cluster compared with healthy controls. **C:** Box plot illustrating the distribution of clinical features across MDS patient clusters. *P* values were calculated using ANOVA with a post hoc Tukey HSD test. *, *P* ≤ 0.05; **, *P* ≤ 0.01.

To understand the distinct epigenetic deregulation associated with MDS patient cluster VI, we annotated the DMRs identified (**Figure 4A**) to neighboring coding and non-coding transcripts. Pathway analysis revealed they targeted genes involved in IL-2 and IL-6 signaling and other inflammation-related pathways (**Figure 4B; Supplemental Table 7)**. Comparing the transcriptional profiles between controls and cluster VI patients, we identified 930 down-regulated and 1,253 up-regulated DEGs (**Supplemental Table 8**). GSEA revealed that while a few of the same inflammation-related pathways seen in the entire cohort of MDS cases (**Supplemental Figure 1C, Supplemental Table 3**) were also upregulated in cluster VI, this cluster was characterized by a less inflammatory profile overall, showing instead distinct enrichment of pathways involved in DNA replication, cell cycle, DNA repair (mismatch repair, base excision repair, and homologous recombination), cysteine and methionine metabolism, and phosphatidylinositol signaling **(Figure 4C; Supplemental Table 7)**.

**Figure 4.**
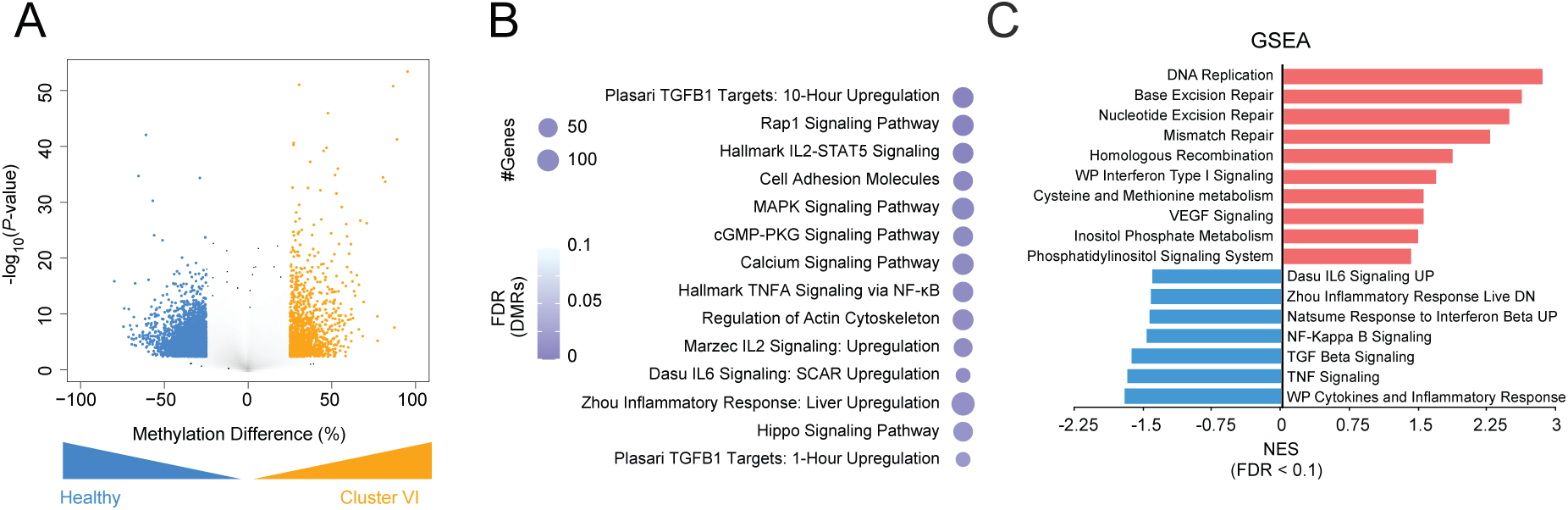
Cluster VI patients display mild epigenetic deregulation and shared clinical and laboratory features. **A:** Volcano plot illustrating methylation difference versus the -log_10_(*P*-value) between healthy controls and MDS Cluster VI. Significant hypomethylated (blue) and hypermethylated (orange) regions in MDS Cluster VI are highlighted (absolute methylation difference ≥ 25%, FDR ≤ 0.05). **B:** Bubble plot depicting enriched pathways (FDR ≤ 0.1) associated with differentially methylated regions in MDS Cluster VI. Bubble size corresponds to the number of genes within each gene set, while color indicates statistical significance. **C:** Bar plot showing the normalized enrichment score (NES) for pathways enriched in MDS Cluster VI compared with healthy controls. Pathways with increased activity are shown in red, while those with decreased activity are shown in blue (FDR ≤ 0.1).

These findings demonstrate how epigenetic profiles reveal novel biology in MDS. Namely, identified a distinct subgroup of MDS patients, defined by a unique DNAme signature and characterized by shared clinical and laboratory features as well as upregulation of cell cycle, DNA repair, and methionine pathways. This may help explain both the paucity of mutations in this group and its less aberrant DNAme profile, as well as an increased rate of incorporation of the nucleoside analog into the DNA, resulting in a more benign clinical presentation and improved response to DNMTi.

### Sensitivity to AZA is associated with specific epigenetic profiles at diagnosis

Using International Working Group criteria^39^, we classified patients as responders (if they achieved a hematologic improvement, partial or complete response within 6 months of treatment initiation), and non-responders (those with stable or progressive disease). PC analysis based on gene expression profiles could not effectively differentiate between AZA responders and non-responders **(Figure 5A)**, while a supervised analysis approach identified only 25 DEGs between the two groups **(Figure 5B; Supplemental Table 9).** Despite this, GSEA revealed distinct pathway enrichment, with sensitivity to DNMTi associated with an expression profile characterized by a lower degree of inflammation (**Figure 5C; Supplemental Table 10**). By contrast, while unsupervised analysis of DNAme profiles also failed to robustly segregate overall AZA responders or even best- responders (i.e. those achieving a complete response) from non-responders **(Supplemental Figure 5A-B**), supervised analysis revealed marked DNAme differences associated with response to AZA, consisting of 1,505 hypomethylated and 1,067 hypermethylated DMRs (absolute methylation difference ≥25% and FDR≤0.05; **Figure 5D; Supplemental Table 11)**. These DMRs were enriched at intergenic regions (DMRs 48% vs. 36% p-value≤0.001), and strongly depleted from promoter regions (DMRs 5% vs. 21%, p-value≤0.001) and CpG islands (DMRs 7% vs. 25%, p-value≤0.001), highlighting a potential role for distal regulatory regions in helping determine response to AZA **(Supplemental Figure 5C)**. When annotated to neighboring genes, like gene expression, these DMRs were enriched in gene sets associated with cell adhesion and TGFb responses **(Supplemental Figure 5D; Supplemental Table 10).**

**Figure 5.**
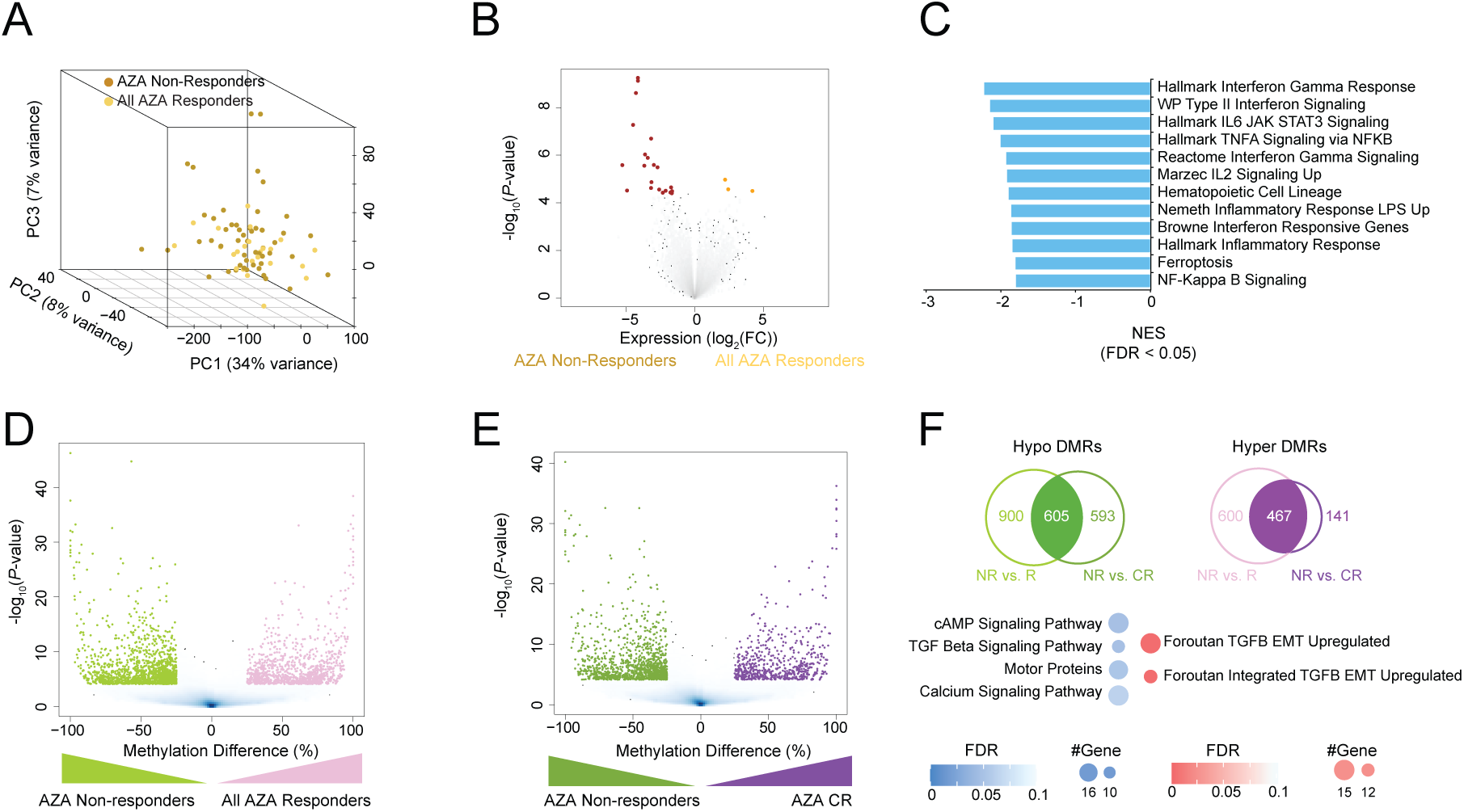
Epigenetic differences distinguish AZA Responders and Non-Responders prior to treatment. **A:** Principal Components Analysis of RNA-seq data from AZA non-responders (brown) and responders (orange). **B:** Volcano plot of the log_2_fold-change (AZA responders/non-responders) gene expression versus the -log_10_(*P*-value). Significant downregulated and upregulated genes in AZA responders are shown in brown and orange, respectively (absolute fold change ≥ 1.5, FDR ≤ 0.05). **C:** Bar plot displaying the normalized enrichment score (NES) for pathways differentially regulated in AZA responders versus AZA non-responders. Upregulated pathways are shown in red, and downregulated pathways are shown in blue (FDR ≤ 0.05). **D:** Volcano plot showing methylation difference in methylated regions versus the -log_10_(*P*- value) between AZA responders and AZA non-responders. Significant hypomethylated and hypermethylated regions in AZA responders are colored in green and light purple, respectively (absolute methylation difference ≥ 25%, FDR ≤ 0.05). **E:** Volcano plot depicting methylation difference versus the -log_10_(*P*-value) between AZA complete responders and AZA non-responders. Significant hypomethylated (green) and hypermethylated (dark purple) regions in AZA complete responders are highlighted (absolute methylation difference ≥ 25% and FDR ≤ 0.05). **F:** Venn diagram (*top*) illustrating the overlap of differentially methylated regions between AZA non-responders vs. responders and AZA non-responders vs. complete responders. The bottom panel shows a bubble plot of enriched pathways (FDR ≤ 0.1) associated with shared hypomethylated and hypermethylated regions. Bubble size corresponds to the number of genes within each gene set, while color indicates statistical significance.

To further refine the epigenetic profiles associated with sensitivity to AZA, we repeated the analysis, restricting the AZA responders’ group to those who achieved complete remission (CR) and compared them to non-responders. This analysis identified 1,198 hypomethylated and 608 hypermethylated DMRs (absolute methylation difference ≥ 25% and FDR ≤ 0.05) associated with achieving a CR **(Figure 5E; Supplemental Table 12)**. Once again, DMRs were enriched at intergenic regions (DMRs 47% vs. 35%, p-value≤0.001), and were strongly depleted from promoter regions (DMRs 6% vs. 22%, p-value≤0.001), and CpG islands (DMRs 9% vs. 26%, p-value≤0.001) **(Supplemental Figure 5E)**. Pathway analysis revealed that while these DMRs were enriched in inflammation pathways as before, they were also enriched in signaling pathways such as cAMP, RAS, calcium signaling, and cell adhesion pathways **(Supplemental Figure 5F; Supplemental Table 13)**. Notably, a subset of DMRs overlapped between the two comparisons, and these overlapping DMRs were enriched in signaling pathways such as cAMP, TGFb, and calcium signaling pathways, indicating that these may play a central role in response to AZA (**Figure 5F; Supplemental Table 14**).

In summary, comprehensive DNAme analysis provided valuable insights into the response of MDS patients to AZA treatment. Moreover, the genomic distribution pattern of these response-associated DMRs explains why promoter-centric analyses did not identify any correlation between DNAme profiles and response to DNMTi and underscores the biological importance of epigenetic deregulation at distal genomic elements.

### DNAme profiles can be harnessed for the robust prediction of therapeutic response

Given the existence of baseline DNAme differences in responders and non-responders, we hypothesized that these unique methylation profiles could be harnessed to predict which patients would be sensitive or resistant to AZA treatment. To test this, we first used stratified randomization to divide patients into a training and independent validation cohort while ensuring a balanced inclusion of responders and non-responders in both groups. The training and independent validation cohorts consisted of 63 and 27 patients, respectively. To avoid any potential overfitting of the model, percent cytosine methylation at 154 DMRs identified between responders and non-responders using only the 63 randomly selected patients in the training cohort was used as input (**Supplemental Table 15**) for a machine-learning approach using random forest (RF) to build a classifier. Five-fold cross-validation was performed, and input features were ranked and selected based on their importance. Ninety-eight selected features were then used to train the final RF model with the best parameters and applied to the 27 patients in the independent blinded test set. The reported area under the receiver operating characteristic curve (ROC-AUC) on the test set was used to evaluate the classifier’s performance (**Figure 6A**). This DNAme RF classifier demonstrated the ability to predict the response to AZA with an AUC score of 0.82, with a sensitivity of 0.7 and a specificity of 0.77 **(Figure 6B**).

**Figure 6.**
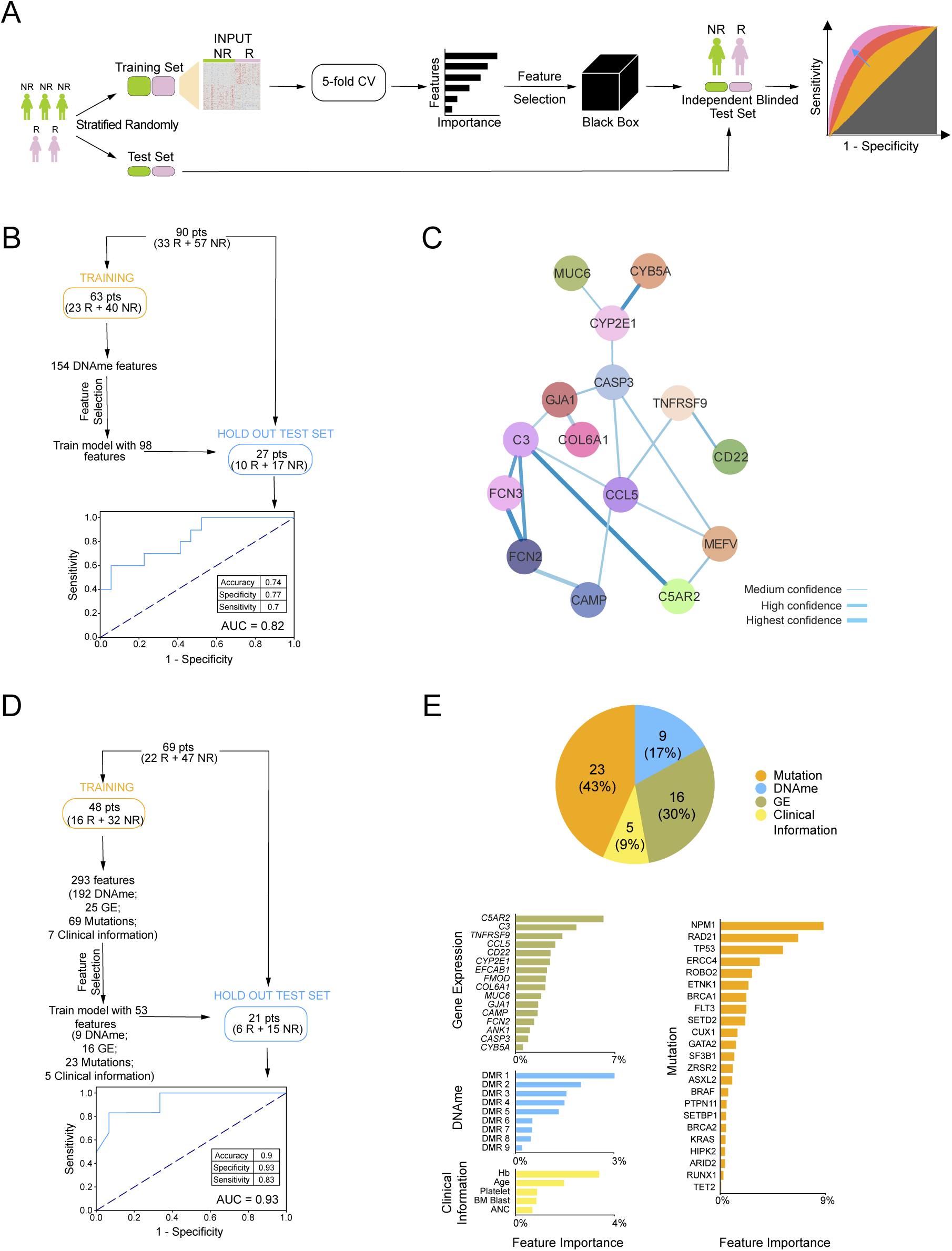
Machine learning models predict response to AZA at baseline. **A:** Schematic representation of the machine learning model used to identify predictive features for AZA response at diagnosis. **B:** Workflow of machine learning predictor utilizing DNA methylation features to distinguish AZA responders from non-responders. The receiver operating characteristic curve of biomarker performance is shown. **C:** Protein-protein interaction network of genes associated with differentially methylated regions within the same topologically associating domain. Line thickness represents connection confidence levels. **D:** Workflow of an integrated machine learning predictor incorporating DNA methylation, gene expression, mutations, and clinical features. The receiver operating characteristic curve of biomarker performance is shown. **E:** Pie chart illustrating the distribution of selected features from the model in D (*top*), and bar plot depicting the importance of each selected feature (*bottom*).

Despite this robustness, we hypothesized that incorporating clinical, mutation, and gene expression information would further improve its performance. Since previous results showed only a few genes with differential expression between responders and non-responders, we first sought to identify genes whose expression is likely epigenetically regulated by DNAme. We previously reported that analyzing gene expression-methylation correlations within functional genomic units determined by topologically associating domains (TADs) leads to the identification of genes regulated via DNAme.^40^ Therefore, we used 2,711 TAD regions identified in CD34+ HSPC and, for each sample for which we had gene expression, mutation, clinical information, and DNAme data available (n=69), we calculated the Pearson correlation between the DNAme level of all the DMRs between responders and non-responders and the gene expression value of each gene transcript within the same TAD. As a result, 4,612 genes were found to be highly correlated with 1,092 DMRs within 820 TADs, with an absolute r≥0.7. Next, we performed network analysis to identify relevant networks of epigenetically regulated genes. The top network identified consisted of 15 genes forming a highly interconnected network with CCL5 as the hub node **(Figure 6C)**. Notably, several genes in this network were regulated via DMRs located at long distances from their loci, again underscoring the importance that distal events play on epigenetic deregulation in MDS **(Supplemental Figure 6A).**

Using this cohort of patients with available DNAme, RNA-seq, mutation, and clinical information, we randomly assigned patients to a training cohort (n=48) and an independent validation cohort (n=21). Using DMRs (n=192), the expression from our CCL5 network (n=25), mutation data (n=69 genes), and clinical parameters (n=7) previously shown to be relevant to MDS biology and prognosis^41–45^ (**Supplemental Table 16**), we performed feature selection as before. Fifty-three features were selected, consisting of a combination of clinical, mutation, expression, and DNAme data. Among the clinical data, hemoglobin, age, platelet count, BM blast percentage, and absolute neutrophil counts (ANC) were included in the final model. Twenty-three gene mutations were included in the RF model. While some of them have been shown to have prognostic impact in MDS (e.g. NPM1^46^, TP53^47^, FLT3^48,49^, and CUX1^50,51^), most of them have not been previously implicated as either prognostic or predictive of response to therapy in MDS. Notably, TET2 mutations, for which conflicting findings regarding their correlation with AZA response have been reported^52–54^, had the lowest feature importance in our model, while ASXL1 and DNMT3A mutations, which have also been reported to correlate with response^55–57^, were not selected at all. When applied to the 21 patients in the blinded independent test set, this new prediction model resulted in a higher AUC score of 0.93, with a sensitivity of 0.83 and specificity of 0.93, demonstrating the increased robustness of this integrative classifier for predicting AZA response **(Figure 6D-E)**.

To determine the relative importance of the different types of features included, we also tested predictive models, excluding each of the different data types one at a time. Exclusion of the DNAme data resulted in the classifier’s lowest performance, with an AUC score of 0.54 (**Supplemental Figure 6B**). Conversely, including DNAme while excluding gene expression features from the same model improved performance to an AUC score of 0.79, with a sensitivity of 0.8 and a specificity of 0.83. However, this was still significantly lower than the complete classifier’s performance (**Supplemental Figure 6C**). Exclusion of either mutation or clinical data also resulted in a decrease in the classifier’s performance, with AUC scores of 0.78 and 0.79, respectively **(Supplemental Figure 6D- E).**

These findings indicate that DNAme data significantly contribute to the predictive power of classifier models, even to a greater extent than gene expression, mutation, and clinical and laboratory data. It is important to note that the highest-performing model resulted when integrating all four types of information, underscoring the synergism among the different features.

## DISCUSSION

The slow DNMTi response and the necessity for ongoing treatment present significant challenges in assessing treatment effectiveness and adjusting therapeutic regimens promptly. By integrating genomic, epigenomic, and transcriptomic data, we gained insights into the complex biological networks underlying MDS heterogeneity, identifying those likely to respond to AZA. Understanding critical genomic regions, gene expression patterns, mutations, and clinical parameters related to AZA sensitivity is essential for developing more targeted and effective therapies. Our study offers initial insights into long-range epigenetic gene regulation mechanisms in MDS pathogenesis. We found that genomic regions affected by DNAme primarily reside in distal regulatory areas. Hi-C sequencing revealed how DMRs control the expression of distal genes within the same TADs, confirming previous findings in AML^40^ and emphasizing the need to focus on promoter and distal regulatory elements in epigenetic dysregulation research in MDS.

Recent efforts have been made to classify MDS patients on the basis of its molecular make-up, resulting in the creation of the International Consensus Classification (ICC)^58^ and the 5th edition of the WHO Classification^59^, and the IPSS-M for prognostication. However, these classifications overlook gene expression and epigenetic information, crucial for understanding the disease. The epigenetic classification in this study identified a novel MDS subgroup (Cluster VI), characterized by a unique phenotype, few mutations, near-normal hemoglobin, lower-risk IPSS-M, and a higher proportion of responders to AZA treatment. Though it shares features with the CCUS-like group described by Bernard et al., that group included both MDS and CMML patients, which are distinct disorders with different molecular profiles and therapy responses. Furthermore, DDX41 mutations, linked to AML and MDS predisposition and favorable outcomes^60^, were more prevalent in this cluster, contrasting with the Bernard classification that separates DDX41 mutation patients despite their shared features with others, as demonstrated in our study.

While prior studies have linked specific mutations to an increased response rate to DNMTi’s, this did not consistently correlate with improved survival, nor were they reproducible across other studies.^52,61,62,53,63–65^ We report the development of robust predictive classifiers by integrating DNAme, gene expression, mutations, and clinical and laboratory parameters. This approach was critical in enhancing the predictive power of our model. Specifically, mutations in 23 genes -including *NPM1, RAD21,* and *TP53*- along with the expression patterns of 16 genes, such as *C5AR2*, *C3,* and *TNFRSF9*, significantly contributed to the model’s performance. While some of these mutations and gene expression patterns have been reported in previous studies on myeloid disorders^66–69^, they have not been collectively used as biomarkers for AZA response. Notably, excluding DNAme from the model resulted in a significant decline in performance, highlighting the importance of DNAme information in capturing disease biology and improving prediction accuracy. Our findings highlight the potential of epigenetic data as predictive biomarkers and emphasize the importance of integrating multiple molecular features for improved predictive accuracy. Prospective validation in larger cohorts will be necessary to fully validate these classifiers for their incorporation into clinical practice.

In conclusion, our integrative approach sheds light on the complexity of MDS and underscores how this molecular interplay influences treatment response. This holistic perspective can offer valuable insights into the molecular mechanisms underlying AZA response in MDS and may ultimately lead to more effective and personalized treatment strategies. Further exploration of the roles of biological pathways identified will be critical for advancing our understanding of MDS pathogenesis and therapeutic approaches.

## ACKNOWLEDGMENTS

We thank all members of the Figueroa Lab for thoughtful discussions and suggestions. This study was supported by award R01HL126947 from the NIH NHLBI, a Leukemia and Lymphoma Society (LLS) scholar award (13-5719), and an LLS SCOR award (7028-22) to MEF. Research reported in this publication was performed in part at the Onco- Genomics Shared Resource (OGSR) of the Sylvester Comprehensive Cancer Center at the University of Miami, RRID: SCR_022502, which is supported by the National Cancer Institute (NCI) of the National Institutes of Health (NIH) under award number P30CA240139. The content is solely the responsibility of the authors and does not necessarily represent the official views of the NIH. Library construction and sequencing services were provided by the University of Miami’s John P. Hussman Institute for Human Genomics Core Facility (RRID:SCR_017828). Samples for this work were provided by the Roswell Park Comprehensive Cancer Centers Biorepository and Laboratory Services Shared Resource supported by National Cancer Institute (NCI) grant P30CA016056.

## AUTHOR CONTRIBUTIONS

MEF conceived and designed the study and supervised all aspects of the analysis and data interpretation; QY and MTM analyzed genome-wide data; RB analyzed mutational data; MT contributed to clinical data organization and annotation; AB, IC, EC, SH, EAG, MYF, CF, JPR, MJR, AED, MAS, and VS contributed clinically annotated samples; MAS and VS provided critical feedback and contributed to data interpretation; QY and MEF wrote the manuscript with approval from all co-authors.

## Data sharing statement

The ERRBS and bulk RNA-seq data have been deposited in the NCBI’s Gene Expression Omnibus (GEO) database under the accession code GSE281562 (RNA-seq) and GSE281610 (ERRBS). Additional ERRBS data for healthy donors was obtained from GSE52945, samples GSM1536373, GSM1536374 and GSM1536375. For normal RNA data, 8 of them were from publications.^1^

