## Supplementary Methods & Figures for "Unique epigenomic signatures identify biologically significant subtypes of MDS and predict response to azacitidine"

**Human bone marrow samples:**

CD34+ cells were isolated from MNC specimens using the Miltenyi MACS magnetic bead purification kit. Frozen human mononuclear cells (MNC) vials were thawed at 37°C, followed by treatment with 200 μL of DNase-I (Roche) at a concentration of 1 mg/mL for 90 seconds. Subsequently, the cells were gently resuspended in Iscove's Modified Dulbecco's Medium (IMDM) containing 40% fetal bovine serum (FBS) to minimize primary human cell mortality. All centrifugation steps were performed at 1500 rpm. After an 8-minute centrifugation, the cells were resuspended in 15 mL of MACS buffer and then incubated with magnetic beads for 50 minutes at 4°C. Following three consecutive washes with MACS buffer (each for 10 minutes at 4°C), the cells were resuspended in 6 mL of MACS buffer and passed through an AUTOMACS instrument using the POSSELD program. After another 8-minute centrifugation at 4°C, the cells were resuspended in RTL buffer with beta-mercaptoethanol. DNA and RNA were extracted from the CD34+ fraction using the AllPrep Micro kit (QIAGEN), while the CD34+ depleted fraction was processed with the AllPrep Mini kit.

**Mutational sequencing:**

Genomic DNA was employed to create libraries for hybrid capture following the manufacturer's protocol from Agilent. Libraries were assessed and then combined in equimolar amounts, aggregating to 500 mg of DNA for up to 24 samples per reaction. The hybridization process was executed using Agilent Custom SureSelect In Solution Hybrid Capture RNA baits, designed to target 443 kbp of exonic DNA with the aid of 16,890 probes, as detailed in the work of Lindsley and colleagues[1]. Each capture reaction underwent a sequence of washing, amplification, and was sequenced on two lanes of an Illumina HiSeq 2000, generating 100-bp paired-end reads.

**Variant Calling**

The analysis proceeded as follows: Fastq files were aligned to the human genome version hg19 using The Burrows Wheeler Aligner (BWA v0.7.12) MEM module, tailored for paired-end reads[2]. To eliminate redundancy, duplicate reads were identified and removed through Picard tools (V1.91). For variant calling, GATK v3.2 was employed after performing base recalibration. Local realignments for insertions and deletions (indels) were executed using reference variant databases[3]. Somatic variants were identified with LoFreq v2.1.1, considering all variants with a frequency of at least 1%[4]. Further filtering of variants occurred post-variant calling, involving specific parameters: variant frequency below 0.05, read depth at the variant site less than 20, GQ and/or QUAL scores less than 30, and IndelRepeatFilter exceeding 8. Variants exhibiting excessive strand bias and indels with Variant Allele Frequency (VAF) less than 10% adjacent to homopolymer repeats were excluded through manual curation. Filtered variants were initially annotated using ANNOVAR[5]. Variants that were predicted to modify splicing were evaluated as described in the work of Jian and colleagues[6]. Variants located outside the protein-coding regions or splice sites of the genes listed earlier and synonymous variants not predicted to impact splicing were excluded.

To eliminate common polymorphisms, variants with population frequencies equal to or greater than 1% in either the 1,000 Genomes[7] or the Exome Aggregation Consortium (ExAC v.3.1) datasets were likewise removed, unless they were also documented as confirmed somatic mutations in the Catalogue of Somatic Mutations in Cancer (COSMIC) [8]. Remaining variants underwent manual review, and those deemed probable somatic-coding variants were included in subsequent analyses. It's worth noting that the variant calling and interpretation processes were conducted without knowledge of sample identifiers and any associated phenotype information.

**Enhanced reduced representation bisulfite sequencing:**

ERRBS was carried out according to established protocols[9], with adjustments made to suit the low input of DNA. Before adapter ligation, the methylated adapters were diluted to a concentration of 150 nmol/L. During the gel size selection step, fragments falling within the range of 150 to 450 base pairs were excised. Subsequently, the prepared libraries were subjected to sequencing on an Illumina HiSeq 3000 platform.

**Differentially methylated regions analysis:**

Reads were trimmed using Trim Galore (version 0.6.2)[10] and aligned to a bisulfite-converted human genome (hg19) using Bowtie2[11] and Bismark[12] (versions 2.4.1 and 0.23.1, respectively). After strand collapsing and filtering by coverage (keep loci with reads between 10-400), methylation loci were tiled by window (25bp) and a methylDiff object was created using MethylSig[13] (version 1.5.0) for regions that were present in at least 3 samples per group. Significant differentially methylated regions (DMRs) were identified as those with an absolute mean methylation difference of ≥ 25 and a DMR q-value of ≤ 0.05. DMRs were then annotated to hg19 using specific parameters: DMRs were assigned to the gene they overlapped. For intergenic DMRs, we assigned them to all neighboring genes within 50 kb. If there was no gene present within a 50-kb window, we assigned the DMR to the nearest transcription start site. We then performed functional annotation with all called methylation tiles as background using ChiPseeker[14] version 1.28.3. Heatmaps were generated with pheatmap[15] 1.0.12, where percent methylation of each DMR for each sample was calculated and plotted using the complete clustering method with Euclidian distance. NA values were depicted in light grey.

**RNA sequencing**

RNA extracted from the CD34+ fraction was employed for RNA-seq. RNA quality was evaluated using an Agilent Tapestation to guarantee high RNA quality. The RNA was treated with Illumina Ribo-Zero. For library preparation, we used the Nugen Ovation Solo RNA-seq ultra-low input kit, following the manufacturer's instructions. Subsequently, the prepared libraries were sequenced on an Illumina NextSeq platform. These libraries were paired-end and they were strand-specific. To normalize the data across runs, ERCC spike-ins were introduced.

**RNA-seq alignment:**

The quality of the resulting reads was assessed using FastQC[16] (v0.11.8) and multiQC[17] (1.11). Then all reads were processed using Trim Galore[10] (version 0.6.2) to trim to paired end 50 basepairs and remove adapters. The resulting reads were aligned to the hg19 Gencode v19 reference genome using the STAR[18] aligner (version 2.7.6a) with specific parameters: outFilterType=BySJout, outFilterMultimapNmax=20, alignSJoverhangMin=8, alignSJDBoverhangMin=1, outFilterMismatchNmax=999, alignIntronMin=20, alignIntronMax=1000000, alignMatesGapMax=1000000, and alignEndsType=EndToEnd.

**Differential gene expression analysis:**
Gene expression was quantified using featureCounts[19] (v2.0.1), which was run in paired-end mode with the hg19 gencode annotation file (excluding entries for ribosomal RNA). Differential expression analysis was performed using DESeq2[20] (v1.32.0) with a multifactor design that accounted for both the donor's condition and any batch effects. Dispersion was calculated across both groups, and pairwise contrasts were established. Genes with an absolute fold change of at least 1.5 and a p-value adjusted for multiple comparisons of less than or equal to 0.05 were considered significant.

**MDS Classification:**
The MDS patient cohort was annotated using the International Prognostic Scoring System (IPSS)[21], the Molecular International Prognostic Scoring System[22], the Revised IPSS (IPSS-R)[23], and the 5th edition of the Classification[24]. For example, under WHO 2022 criteria, MDS-bi *TP53* includes cases with either two or more *TP53* mutations, or one *TP53* mutation accompanied by a complex karyotype involving the chromosome on which *TP53* is located.

**GSEA:**
Using a pre-ranked list of genes based on the Wald statistic from the DESeq2[20] output, Gene Set Enrichment Analysis (GSEA)[25] (v4.2.3) was conducted. The gene set size was limited to 15-500 genes, and a weighted enrichment score was utilized.

**DNA methylation to gene expression correlation analysis:**
Hi-C sequencing was used to analyze CD34+ HSPCs, and 2,711 topologically associating domain (TAD) regions were identified using previously established methods, as described in the reference study[26]. Differentially methylated regions that exhibited coverage between 10-400 in more than 5 samples for both AZA responders and non-responders were chosen for conducting Pearson correlation analysis. The correlation

analysis involved examining the transcripts per million (TPM) values of all expressed genes within the same TAD region as the selected DMRs. Highly correlated pairs of DMRs and genes with absolute correlation coefficient ≥ 0.7 were identified.

**Protein-protein interaction:**

The genes highly correlated with DMRs were analyzed using the STRING[27] database (v11.5) to construct protein-protein interaction networks. Then network topology measures such as degree centrality and betweenness centrality were calculated using the NetworkAnalyzer plugin in Cytoscape[28] (v3.9.1) to identify highly connected and influential nodes in the network. The top sub-networks with a confidence score of at least 0.4 (medium confidence) were extracted and visualized using Cytoscape. The nodes were colored and made transparent based on their bundling strength, and the thickness of the edges represented the confidence of the correlations between genes. The identified nodes were further analyzed for their functional annotations and pathways using Kyoto Encyclopedia of Genes and Genomes (KEGG)[29] pathway enrichment analyses.

**Machine learning classifier:**

The samples were randomly stratified and divided into training and testing cohorts, with one-third of the samples reserved for testing purposes using train_test_split() function from sklearn[30] (v.1.2.2). Differentially methylated regions and differentially expressed genes between responders and non-responders in the training cohort were ascertained using previously described methodologies, and their DNA methylation levels (without missing values) and/or gene expression levels, mutation status, and other clinical parameters were extracted for both the training and testing cohorts. The resulting matrices were standardized using the StandardScaler() function to ensure comparability of means and standard deviations.

For feature selection, Recursive feature elimination, L1-based feature selection and mutual information based techniques were employed. GridSearchCV() with 5-fold cross validation was performed to optimize model hyperparameters.

With the optimal parameters from the training cohort, the random forest algorithm was utilized to predict response employing the RandomForestClassifier() function from sklearn.ensemble and the area under the receiver operating characteristic curve score was computed.

**SUPPLEMENTARY TABLES:**

**Supplemental Table 1: Cohort Characteristics.** A table describing the patient information for samples used in the study.

**Supplemental Table 2: Differentially expressed genes in MDS vs. Healthy.** Related to Supplementary Figure 1. List of differentially expressed genes in MDS compared to healthy controls (absolute fold change ≥ 1.5, FDR ≤ 0.05). A positive log2 fold-change corresponds to increased gene expression in MDS patients.

**Supplemental Table 3: Pathway analysis of differentially expressed genes and differentially methylated regions in MDS vs. Healthy.** Related to Figure 1 and Supplementary Figure 1. Enriched pathways associated with differentially methylated regions between MDS and healthy controls, as well as enriched pathways identified through gene set enrichment analysis using RNA-seq data.

**Supplemental Table 4: Differentially methylated regions of MDS vs. Healthy.** Related to Figure 1. List of differentially methylated regions in MDS compared to healthy controls (absolute methylation difference ≥ 25%, FDR ≤ 0.05).

**Supplemental Table 5: Clincal and molecular characteristics of MDS DNA methylation clusters.** Related to Figure 2. Significant molecular and clinical features of each MDS cluster, identified using Fisher's exact test

**Supplemental Table 6: Differentially methylated regions of each Cluster vs. Healthy.** Related to Figure 3, 4 and Supplementary Figure 4. List of differentially methylated regions in each MDS cluster compared to healthy controls (absolute methylation difference ≥ 25%, FDR ≤ 0.05).

**Supplemental Table 7: Pathway analysis of differentially expressed genes and differentially methylated regions in Cluster VI vs. Healthy.** Related to Figure 4. Enriched pathways associated with differentially methylated regions in MDS Cluster VI compared to healthy control, as well as enriched pathways identified through gene set enrichment analysis using RNA-seq data.

**Supplemental Table 8: Differentially expressed genes of Cluster VI vs. Healthy**. Related to Figure 4. List of differentially expressed genes in MDS Cluster VI compared to healthy controls (absolute fold change ≥ 1.5, FDR ≤ 0.05). A positive log2 fold-change corresponds to increased gene expression in MDS Cluster VI.

**Supplemental Table 9**: **Differentially expressed genes of AZA responders vs. Non-responders.** Related to Figure 5. List of differentially expressed genes in AZA responders compared to AZA non-responders (absolute fold change ≥ 1.5, FDR ≤ 0.05). A positive log2 fold-change corresponds to increased gene expression in AZA responders.

**Supplemental Table 10: Pathway analysis of differentially expressed genes and differentially methylated regions in of AZA responders vs. Non-responders.** Related to Figure 5 and Supplementary Figure 5. Enriched pathways associated with differentially methylated regions in AZA responders compared to non-responders, as well as enriched pathways identified through gene set enrichment analysis using RNA-seq data.

**Supplemental Table 11: Differentially methylated regions of AZA responders vs. Non-responders.** Related to Figure 5. List of differentially methylated regions in AZA responders compared to AZA non-responders (absolute methylation difference ≥ 25%, FDR ≤ 0.05).

**Supplemental Table 12: Differentially methylated regions of AZA Complete responders vs. Non-responders.** Related to Figure 5. List of differentially methylated regions in AZA complete responders compared to AZA non-responders (absolute methylation difference ≥ 25%, FDR ≤ 0.05).

**Supplemental Table 13: Pathway analysis of differentially expressed genes and differentially methylated regions in of AZA Complete responders vs. Non-responders.** Related to Figure 5 and Supplementary Figure 5. Enriched pathways associated with differentially methylated regions in AZA complete responders compared to non-responders, as well as enriched pathways identified through gene set enrichment analysis using RNA-seq data.

**Supplemental Table 14: Common pathways between AZA R vs. Non-responders and AZA Complete responders vs. Non-responders.** Related to Figure 5. Enriched pathways associated with shared hypomethylated and hypermethylated regions in AZA Responders vs. non-responders and AZA complete responders vs. non-responders.

**Supplemental Table 15: Machine Learning models using DNAme data.** Related to Figure 6. Input data matrix and train/test dataset used for machine learning modeling based on DNA methylation data.

**Supplemental Table 16: Machine Learning models using DNAme, gene expression, mutation and clinical data.** Related to Figure 6. Input data matrix and train/test dataset for machine learning modeling using DNA methylation, gene expression, mutation and clinical data.

### SUPPLEMENTARY FIGURES:

#### Supplemental Figure 1:

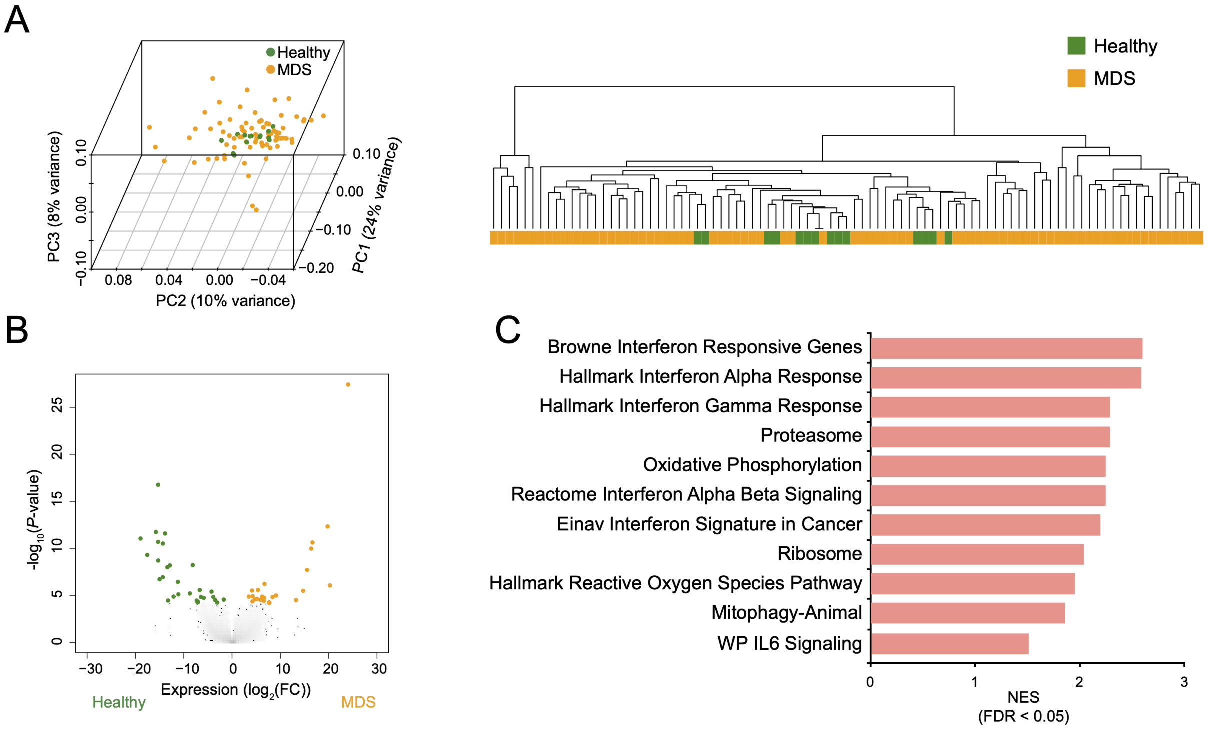


**Supplemental Figure 1. Transcriptomic Profiling in MDS Patients**
**A:** Correspondence analysis (*left*) and hierarchical clustering (*right*) of RNA-seq data, comparing healthy controls (green) and MDS patients (yellow)
**B:** Volcano plot of the $log_2$fold-change (MDS/Healthy) gene expression versus the -$log_{10}$(*P*-value). Downregulated and upregulated genes in MDS patients are shown in green and yellow, respectively (absolute fold change ≥ 1.5, FDR ≤ 0.05).
**C:** Bar plot displaying the normalized enrichment score (NES) for pathways enriched in MDS compared with healthy controls. Pathways with increased activity are shown in red, while those with decreased activity are shown in blue (FDR ≤ 0.05).

**Supplemental Figure 2:**

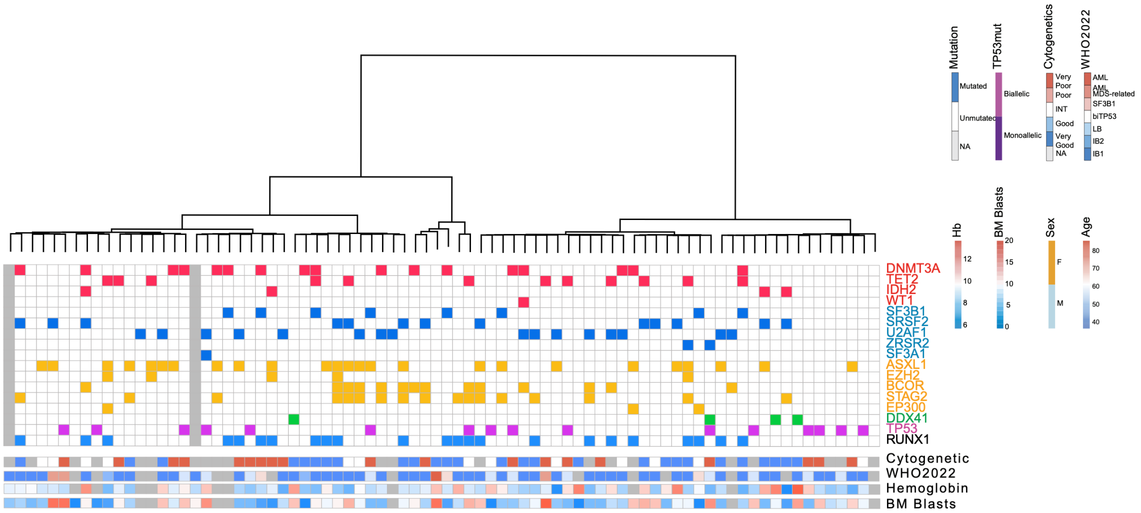


**Supplemental Figure 2. Clustering of MDS Patients Based on Gene Expression Level**

Heatmap and hierarchical clustering of MDS patients based on RNA-seq data, Corresponding mutation and clinical annotations are displayed below, with patients harboring gene mutations color-coded.

**Supplemental Figure 3:**

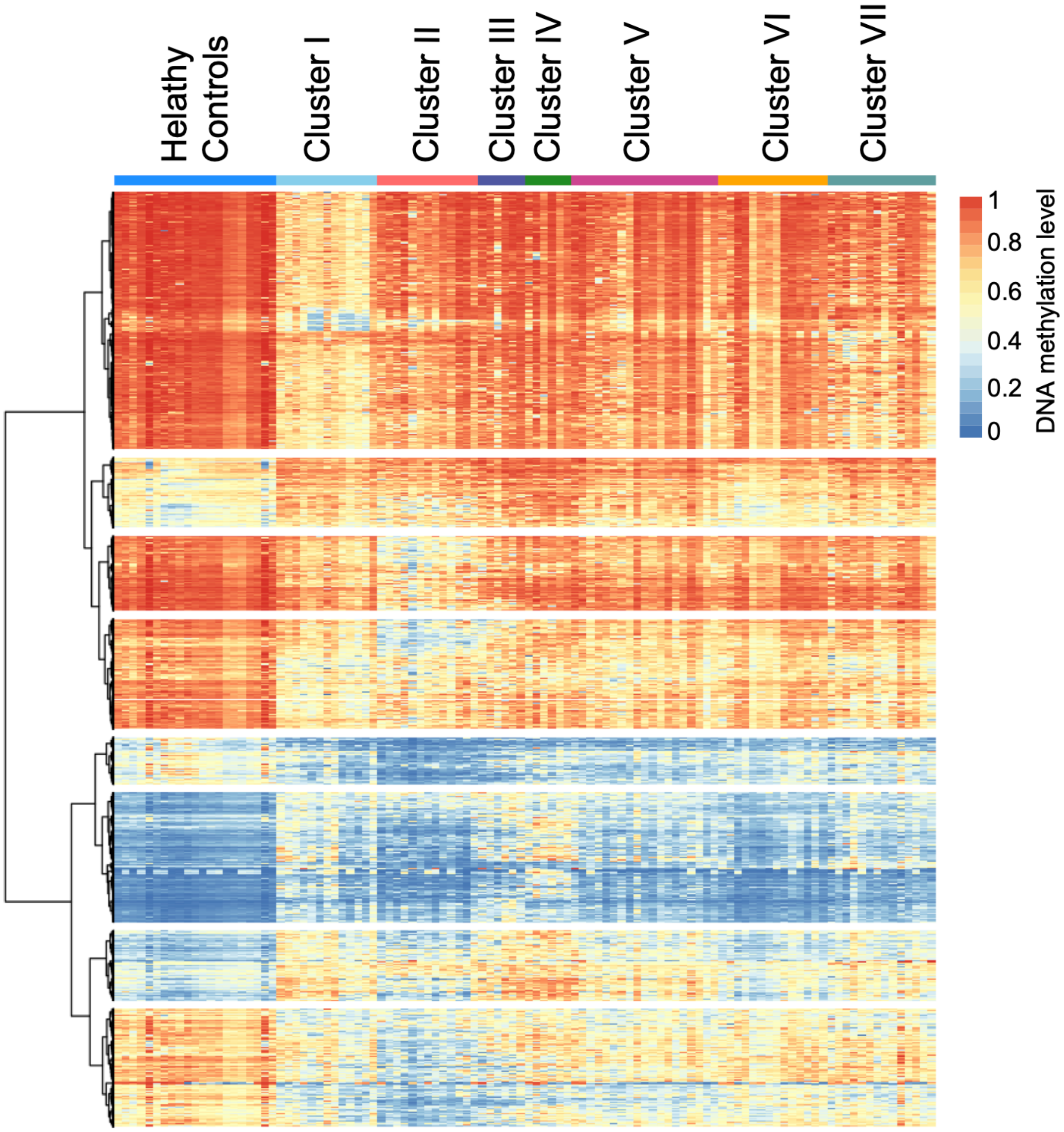


**Supplemental Figure 3. Heatmap of DNA Methylation Levels in MDS patient clusters** Heatmap illustrating DNA methylation levels across seven MDS patient clusters and healthy controls.

**Supplemental Figure 4:**

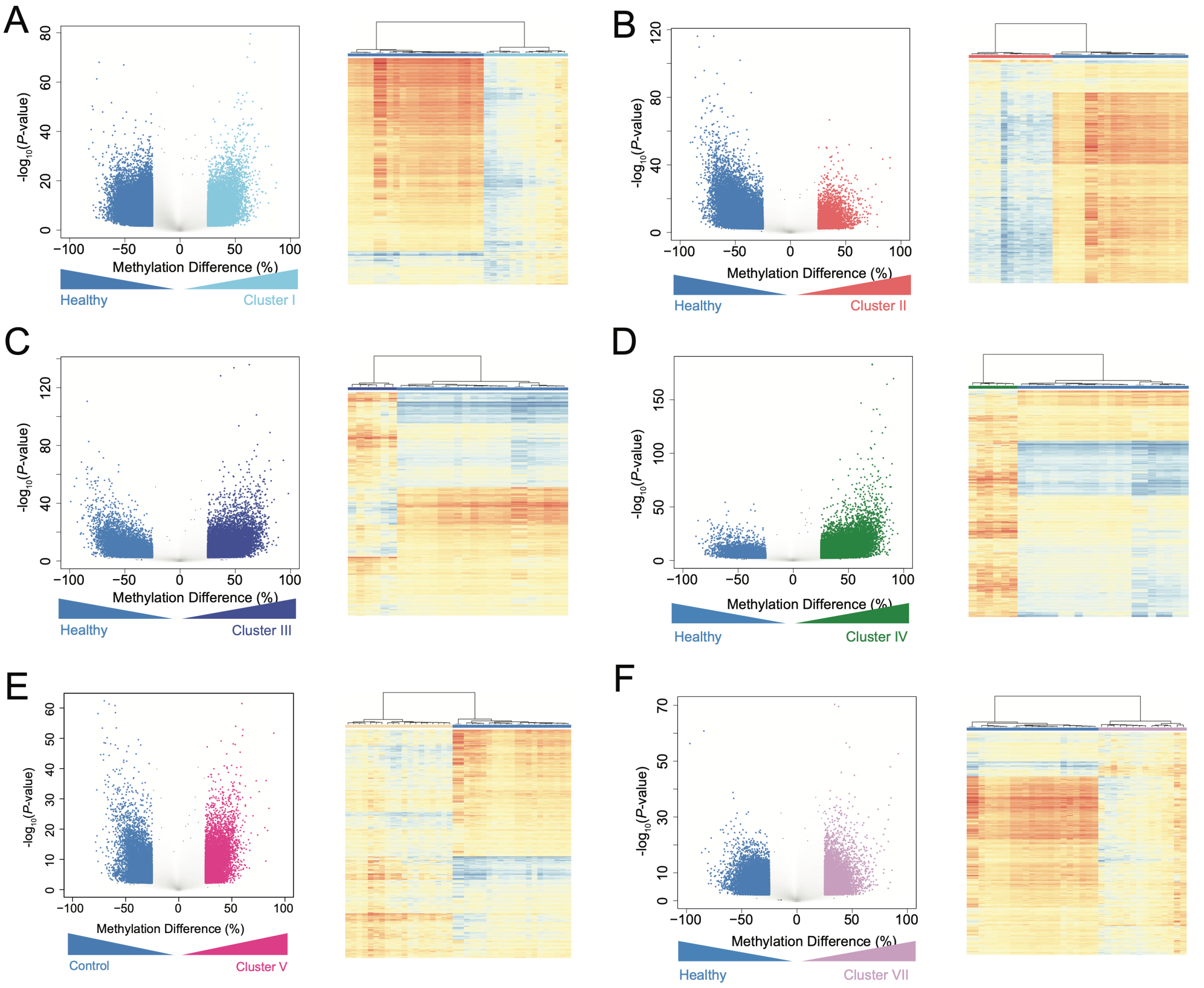


**Supplemental Figure 4. Differential Methylation in MDS Clusters Compared to Healthy Controls**

**A-F:** Volcano plots (*left*) illustrating methylation differences versus the $-\log_{10}$(*P*-value) between healthy controls and each MDS cluster (absolute methylation difference ≥ 25%, FDR ≤ 0.05). Corresponding heatmaps (*right*) display methylation levels of differentially methylated regions in healthy controls (blue bar) and MDS clusters: I (light blue), II (red), III (dark blue), IV (green), V (dark pink), and VII (purple).

**Supplemental Figure 5:**

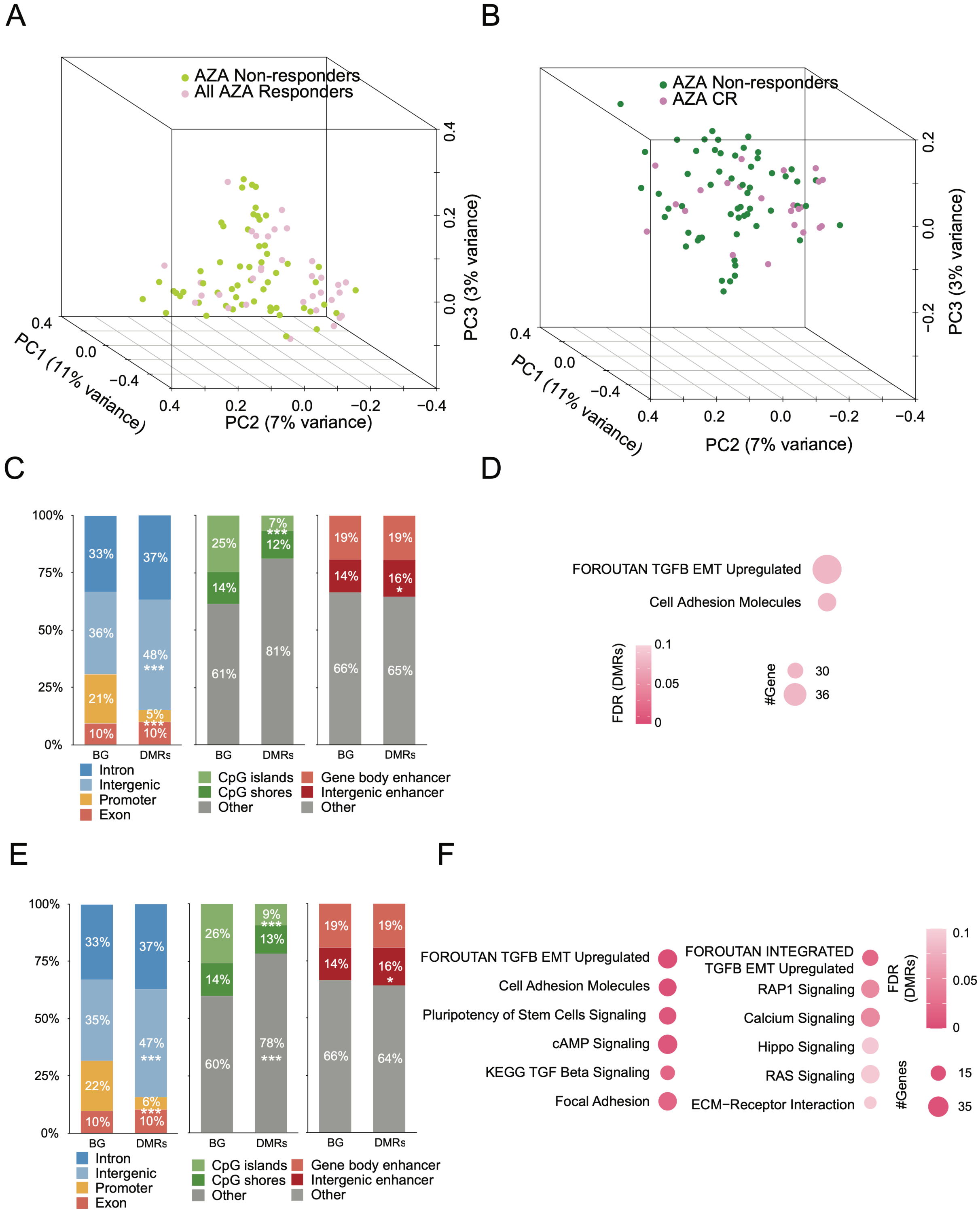


**Supplemental Figure 5. Characterizing Methylation Differences and Pathway Enrichment in AZA Non-Responders and Responders/Complete Responders**

**A:** Principal Component Analysis of ERRBS data comparing AZA non-responders (green) and all responders (light purple).
**B:** Principal Component Analysis of ERRBS data comparing AZA non-responders (dark green) and complete responders (dark purple).
**C:** Stacked bar plot showing the distribution of differentially methylated regions between AZA non-responders and responders across genomic regions, CpG islands, and enhancers. *P* values were calculated using a binomial test. *, $P \leq 0.05$; ***, $P \leq 0.001$.
**D:** Bubble plot representing enriched pathways (FDR ≤ 0.1) associated with differentially methylated regions in AZA responders compared with AZA non-responders, using all CpG tiles as the background. Bubble size corresponds to the number of genes in each pathway, while color indicates statistical significance.
**E:** Stacked bar plot showing the distribution of differentially methylated regions between AZA non-responders and complete responders across genomic regions, CpG islands, and enhancers. *P* values were calculated using a binomial test. *, $P \leq 0.05$; ***, $P \leq 0.001$.
**F:** Bubble plot representing enriched pathways (FDR ≤ 0.1) associated with differentially methylated regions in AZA complete responders compared with AZA non-responders, using all CpG tiles as the background. Bubble size corresponds to the number of genes in each pathway, while color indicates statistical significance.

**Supplemental Figure 6:**

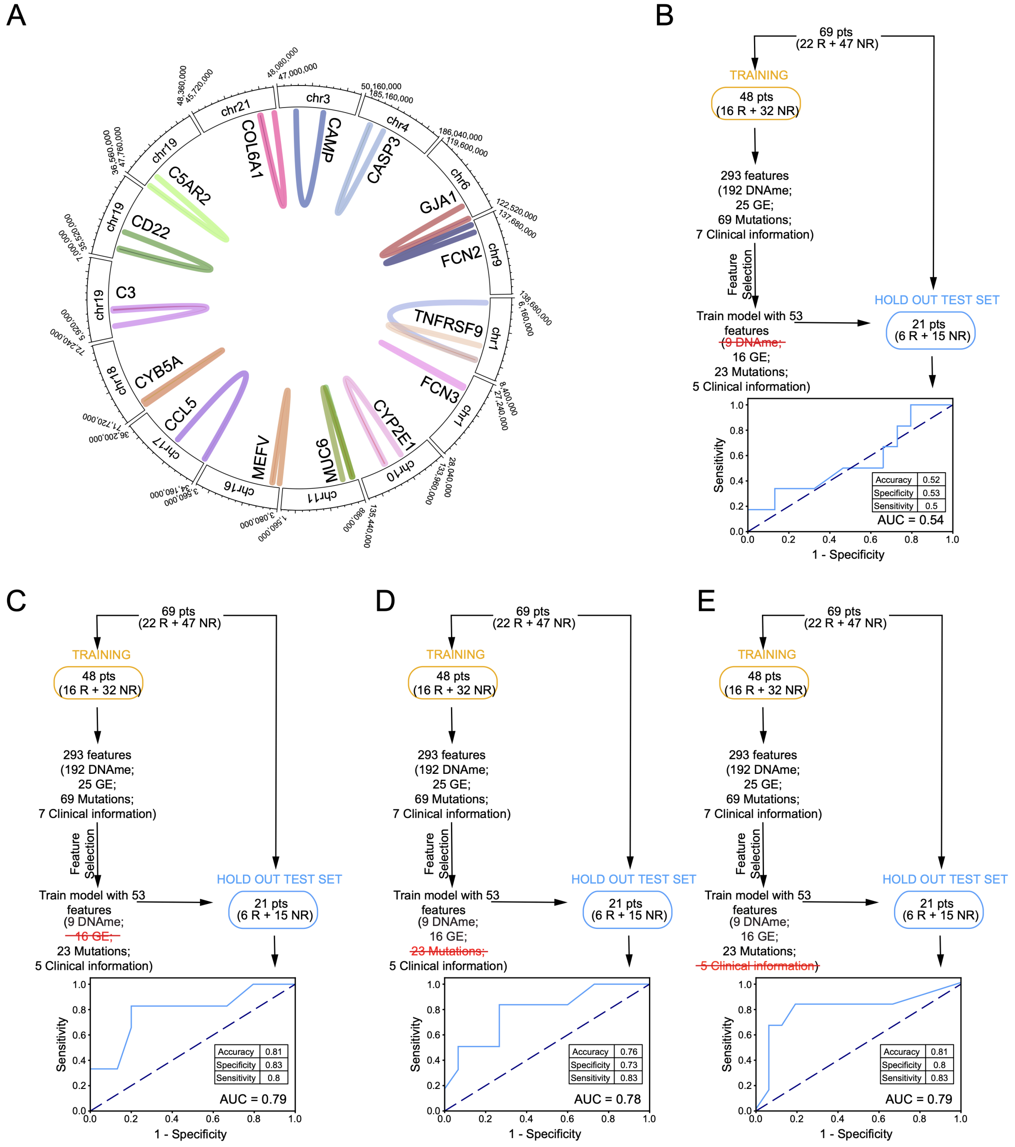


**Supplemental Figure 6. Genes Correlated with DMRs in the Same TAD Regions and Machine Learning Performance with Different Input Features**

**A:** Circos plot illustrating the genomic coordinates of differentially methylated regions and their correlated genes (|Pearson correlation| > 0.7) within the same topologically associating domain regions.

**B:** Workflow of machine learning predictor (from Figure 6D) for distinguishing AZA responders from non-responders after removing DNA Methylation features.

**C:** Workflow of machine learning predictor (from Figure 6D) after removing gene expression features.

**D:** Workflow of machine learning predictor (from Figure 6D) after removing mutation features.

**E:** Workflow of machine learning predictor (from Figure 6D) after removing clinical features.
